# Discovery of novel RNA viruses in commercially relevant seaweeds *Alaria esculenta* and *Saccharina latissima*

**DOI:** 10.1101/2024.05.22.594653

**Authors:** Rob J. Dekker, Wim C. de Leeuw, Marina van Olst, Wim A. Ensink, Selina van Leeuwen, Job Cohen, Klaas R. Timmermans, Timo M. Breit, Martijs J. Jonker

## Abstract

Seaweeds are increasingly recognized as sustainable food sources; however, their large-scale cultivation faces challenges similar to land crops, including susceptibility to pathogens. Plant viruses pose a significant threat to global food security, yet little is known about the diversity of viruses in seaweeds. This study investigates virus-associated small interfering RNA (siRNA) responses in commercially relevant seaweed species to understand RNA virus diversity, particularly in edible varieties. Through small RNA sequencing of 16 samples from *Saccharina latissima* and *Alaria esculenta*, we identified three new RNA viruses Aev-NL1, Slv-NL2 and Slv-NL3, and one new DNA virus (phaeovirus). The partial genome of the new DNA virus was discovered in the *A. esculenta* samples and shared 67% DNA sequence identity with the major coat protein of the large double-stranded DNA phaeovirus *Feldmannia irregularis* virus a. In four out of five *A. esculenta* samples, a new bisegmented ormycovirus-like RNA virus (Aev-NL1) was identified. A similar new virus, Slv-NL1, was found in previously published *S. latissima* RNA-seq data, sharing 87% sequence identity with Aev-NL1. Lastly, two novel RNA viruses, Slv-NL2 and Slv-NL3, were discovered in all eight *S. latissima* samples sharing limited similarity at the genome level but high sequence identity at protein level of both ORFs (>94%). Further investigation of the novel viruses’ presence across our limited set of samples revealed no conclusive associations with diseased seaweed phenotypes. The discovery of four new viruses in only a limited set of samples highlights the presence of previously unrecognized viral diversity in seaweed, thereby underscoring the importance of understanding viral diversity in seaweed as its virome is currently understudied.

## Introduction

Seaweeds, i.e. macroalgae, are gaining traction as sustainable food source (Rogel-Castillo *et al*., 2023). However, with respect to pathogens, large scale production of seaweed is facing the same challenges as commercially grown land crops (Savery *et al*,. 2019). Plant diseases caused by pathogens are worldwide responsible for up to 40% of crop yield losses. Half of these pathogen-caused diseases have a viral origin, which makes plant viruses a severe threat to global food security (Savary *et al*., 2019; He & Creasey Krainer, 2020). However, in contrast to plant viruses, little is known about the diversity of viruses in seaweed, especially RNA viruses (Schroeder and McKeown, 2021). Given the ancient evolutionary origin of seaweeds, as well as the abundant presence of RNA viruses in plants (Wu *et al*., 2022), it is to be expected that seaweeds are host to many RNA virus species. To get a better understanding of seaweed RNA viruses, especially in edible species, we investigated the virus-associated siRNA response in commercially relevant seaweed species with and without unhealthy phenotypes, such as bleaching spots. Although there is limited scientific literature on virus related siRNAs, the evidence for siRNA in algae is mounting (Cerutti & Ibrahim, 2011).

In our study into virus siRNAs in algae, we encountered siRNAs directed at a DNA virus, as well as RNA viruses. In this report we describe the genome sequences of three new RNA viruses, of which two were identified by evaluating the smallRNAs (sRNAs) present in *Saccharina latissima* and *Alaria esculenta* seaweed samples.

## Material and methods

### Samples

We obtained 24 samples: 8 *Ulva lactuca* (green algae) samples (S1-S8), 11 *Saccharina latissima* (brown algae) samples (S9-S16 and S26-S28) of which 7 had a bleaching phenotype (S11-S16, Supplemental Figure S1), and 5 *Alaria esculenta* (brown algae) samples (samples S21-S25), from a Dutch research facility in the south of the Netherlands (NIOZ, S1-S16) and from a commercial development facility in the north-west of the Netherlands (samples S21-S28) (Table 1).

**Table 1.**
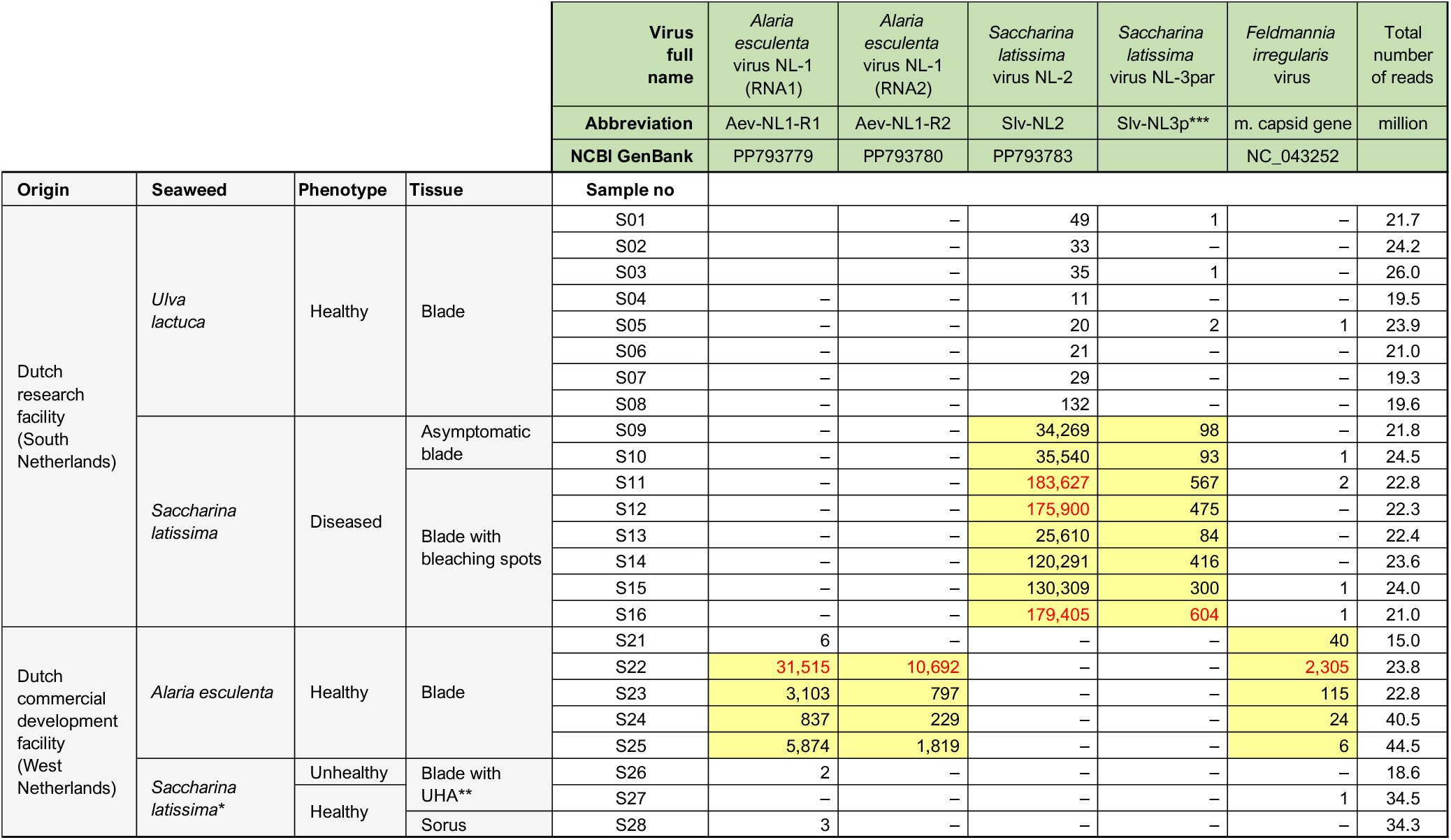
sRNA-seq read counts associated with the new seaweed viruses Aev-NL1, Slv-NL2, Slv-NL3 and Feldmannia irregularis virus. Origin and phenotype description of the samples is shown and the total number of obtained sRNA-seq reads per sample is indicated. *Sample S26 was obtained from the United Kingdom, S27 from Scotland, and S28 from Germany. **UHA is unhealthy appearance of the blades (dark sports). ***Partial genomic sequence known, similar to Slv-NL2, only uniquely-mapping reads counted.

### RNA isolation

RNA was isolated using a CTAB extraction buffer as described previously (Sim *et al*., 2013) with a minor adaptation of method 5, using a final concentration of 2.5 M LiCl. The concentration of the RNA was determined using a NanoDrop ND-2000 (Thermo Fisher Scientific) and the RNA quality was assessed using the 2200 TapeStation System with Agilent RNA ScreenTapes (Agilent Technologies).

### Small RNA-seq (sRNA-seq)

Barcoded sRNA-seq libraries were generated from the total RNA extractions according to the manufacturers’ protocols using the Small RNA-seq Library Prep Kit (Lexogen). The size distribution of the libraries with indexed adapters was assessed using a 2200 TapeStation System with Agilent D1000 ScreenTapes (Agilent Technologies). The sRNA-seq libraries were clustered and sequenced (1 × 75 bp) on a NextSeq 550 Sequencing System (Illumina) using a NextSeq 500/550 High Output Kit v2.5 (75 Cycles) (Illumina).

### Quantitative RT-PCR

Reverse transcription was performed on 300 ng of total RNA primed with a mixture of 2 μM of each respective forward and reverse primer. SuperScript IV Reverse Transcriptase (Thermo Fisher Scientific) was used according to the manufacturer’s instructions with a 1-hour incubation at 55°C. cDNA was diluted 10-fold and subjected to quantitative PCR using the PrimeTime™ Gene Expression Master Mix (Integrated DNA Technologies, Inc.) and QuantStudio 3 Real-Time PCR System (Thermo Fisher Scientific). Primers and probes (Integrated DNA Technologies, Inc. ) specific for Slv NL-2 were CCTATCTGGACCGTCATTTC (forward), AGCCATACAGAC ATATCACATC (reverse), 56-FAM/TGATCTTCT/ZEN/CCGTATTGGAGCTGC/3IABkFQ (probe) and for Slv NL-3 CGAAGGGTCCGAACTTTATC (forward), AAGACCACTCTGGAACTCA (reverse) and 56-FAM/TCCGACGAG/ZEN/ TATTGGTGAAGGAAC/3IABkFQ (probe).

### RACE-sequencing

Total RNA was polyadenylated using *E. coli* Poly(A) Polymerase (New England Biolabs) as per the manufacturer’s guidelines. Subsequently, cDNA synthesis was carried out employing SuperScript IV (Thermo Fisher Scientific) and the oligo(dT)-adapter primer GCGAGCACAGAATTAATACGACTCACTATAGG(T)_30_VN. In order to obtain overlapping halves of the entire Slv-NL2 genomic sequence, including its complete 5’ and 3’ ends, a RACE PCR strategy was used on the cDNA using two primers combinations. For the 5’ half, the Slv-NL2-specific reverse primer TGCCTCGTATCCTCCCAGTCTG was utilized in conjunction with a primer complementary to the adapter GCGAGCACAGAATTAATACGACTC. Likewise, the 3’ half was amplified using the Slv-NL2-specific forward primer TCATGCTTGAAGAGGGCCTG together with the same adapter primer. Size validation of the PCR products was performed using the 2200 TapeStation System with Agilent D1000 ScreenTapes. Subsequent amplicon sequencing was carried out utilizing a Native Barcoding Kit 24 V14 (SQK-NBD114.24) kit together with a Flongle (R10.4.1) flow cell (Oxford Nanopore Technologies) following the manufacturer’s instructions.

### Bioinformatics analyses

Sequencing reads were trimmed using trimmomatic v0.39 (Bolger *et al*., 2014) [parameters: LEADING:3; TRAILING:3; SLIDINGWINDOW:4:15; MINLEN:19]. Mapping of the trimmed reads to the NCBI virus database was performed using Bowtie2 v2.4.1 (Langmead *et al*., 2012). Contigs were assembled using all reads as input with SPAdes De Novo Assembler (Prjibelski *et al*., 2020) with parameter settings: only-assembler mode, coverage depth cutoff 10, and kmer length 17, 18 and 21.

Scanning of contig sequences for potential RdRp-like proteins was performed using PalmScan (Babaian *et al*., 2022) and LucaProt (Hou *et al*., 2023).

## Results

For the discovery of new viruses in seaweed potentially related to disease phenotypes, we isolated and sequenced sRNAs from 8 *U. lactuca*, 11 *S. latissima*, and 5 *A. esculenta* samples with and without unhealthy phenotypes (Table 1; Supplemental Figure S1). sRNA-seq analyses yielded on average 24.6 million reads per sample. By mapping the sRNA-seq reads with reduced stringency to GenBank virus accessions, we were unable to detect the presence of any known virus. Hence, we employed a contig building and similarity comparison strategy to detect potential new viruses. Using this approach we were able to identify the presence of several unknown viruses in *S. latissimi* and *A. esculenta*, but none in *U. lactuca*.

### A new phaeovirus in *Alaria esculenta*

The first putative new virus we detected was represented by two contigs (combined 1,654 bp) obtained from the *A. esculenta* samples that showed 67% DNA sequence identity to the major coat protein of the large dsDNA phaeovirus *Feldmannia irregularis* virus a (FirrV-1,∼180 kb; Delaroque *et al*., 2003; Supplemental File S1), which is known to infect brown algae of the genus *Feldmannia irregularis*. At protein level, the translated contigs showed 66% identity at a 94% coverage (Supplemental File S1). Given that the major coat protein is likely to be highly expressed, it seems logical that the detected sRNAs originate from mRNA of this gene. Given the wide length distribution of mapping sRNA-seq reads (Figure 1C), it can be concluded that the sequenced RNA molecules likely originate from (degraded) mRNA of the coat protein and not siRNAs. Still, virus-related sequencing reads were detected in all five samples from *A. esculenta* blades, albeit with a wide range (Table 1). Unexpectedly, this new phaeovirus in a Laminariales brown alga is most similar to ones found in Ectocarpales brown algae. As all *A. esculenta* samples came from healthy individuals, there seems to be no obvious disease phenotype associated with this new phaeovirus (McKeown, 2019).

**Figure 1.**
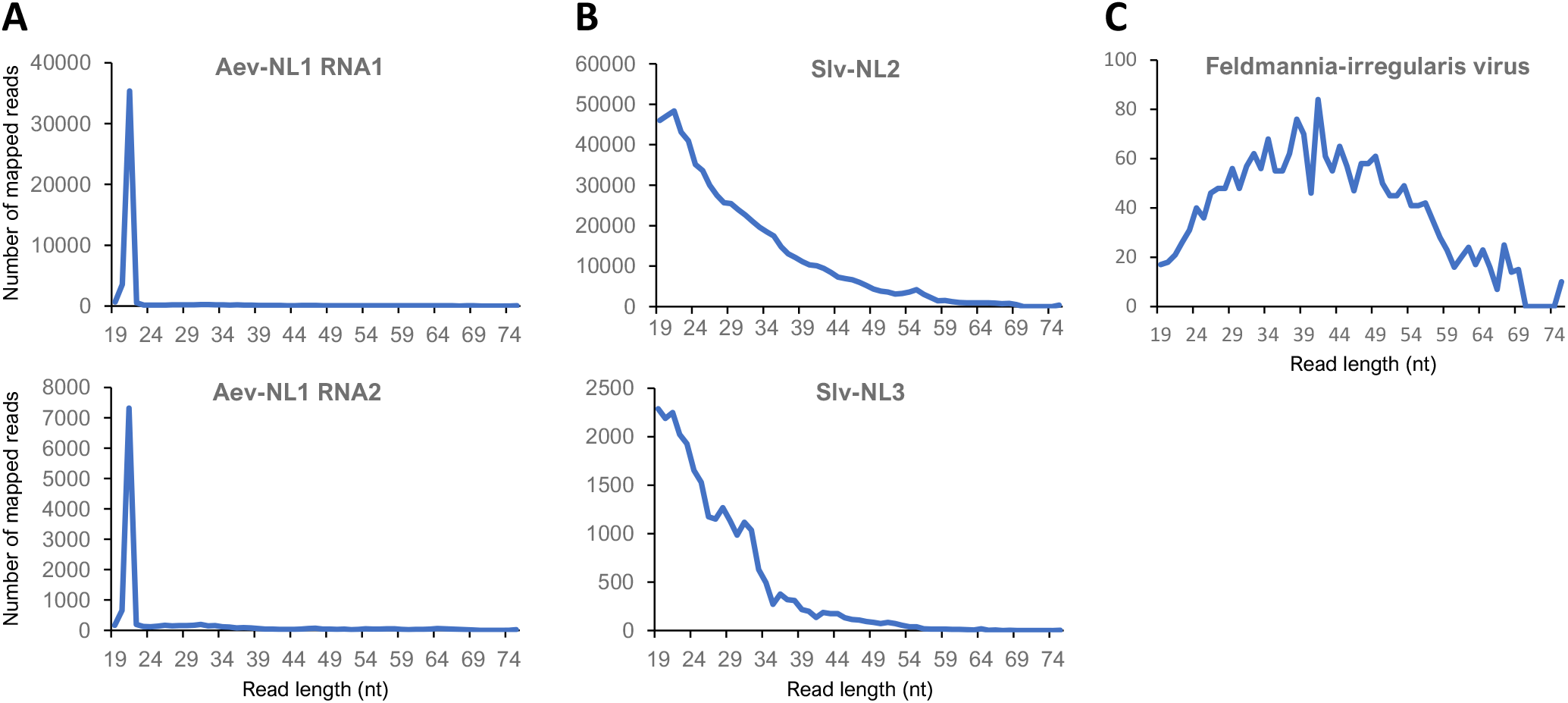
Read length distribution of reads that map to new virus sequences. Length distribution plots of sRNA-seq reads that map to A) Aev-NL2-RNA1 or Aev-NL2-RNA2 genomic sequences; B) Slv-NL2 and Slv-NL3 genomic sequences; C) new Feldmannia-irregularis coat protein genomic sequence.

### A new ormycovirus-like RNA virus in *Alaria esculenta*: Aev-NL1

Using bioinformatics analyses including assembly of all the sRNA-seq reads of the complete experiment into contigs, we discovered a new RNA virus present in four out the five *A. esculenta* samples (Table 1). This new virus, arbitrarily named *Alaria esculenta*-virus-NL1 (Aev-NL1), appears to consist of two RNA molecules; RNA1 (PP793779), 2,420 nt and RNA2 (PP793780), 2,040 nt (Figure 2A; Supplemental File S2). We could not detect any DNA or RNA sequence in the NCBI-GenBank that showed any relevant similarity to the new viral nucleotide sequences. The virus RNAs code for a 781 aa putative RNA-dependent RNA-polymerase (RdRp) protein and a 625 aa putative coat protein, respectively (Figure 2B; Supplemental File S2). At protein level, ORF1-protein from Aev-NL1-RNA1 showed 32% identity at an 80% coverage to the putative RdRp protein from Erysiphe lesion-associated ormycovirus 2 (Forgia *et al*., 2022), whereas ORF2-protein from Aev-NL1-RNA2 showed 36% identity at a 65% coverage to a hypothetical protein from Erysiphe lesion-associated ormycovirus 3 (Figure 2C; Supplemental File S2; Supplemental Table S1). The RdRp classification was confirmed using LucaProt (Hou *et al*., 2023), but no RdRp palmprint barcode could be detected by palmID analysis (Babaian *et al*., 2022). However, additional evidence for the viral origin of the discovered RNA molecules was found in that almost all mapping sRNA-seq reads are 21 nt long (Figure 1A) and have a T nucleotide as 5’ terminus (Table 2), both of which are hallmarks for virus-derived siRNAs.

**Table 2.**
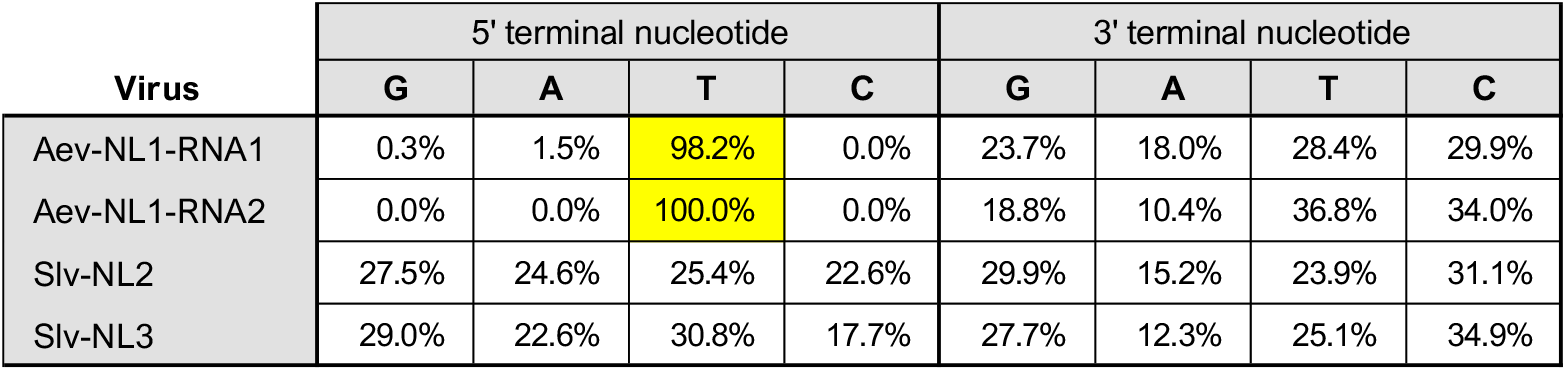
Nucleotide occurrence at 5’ and 3’ terminus of virus associated sRNA-seq reads.

**Figure 2.**
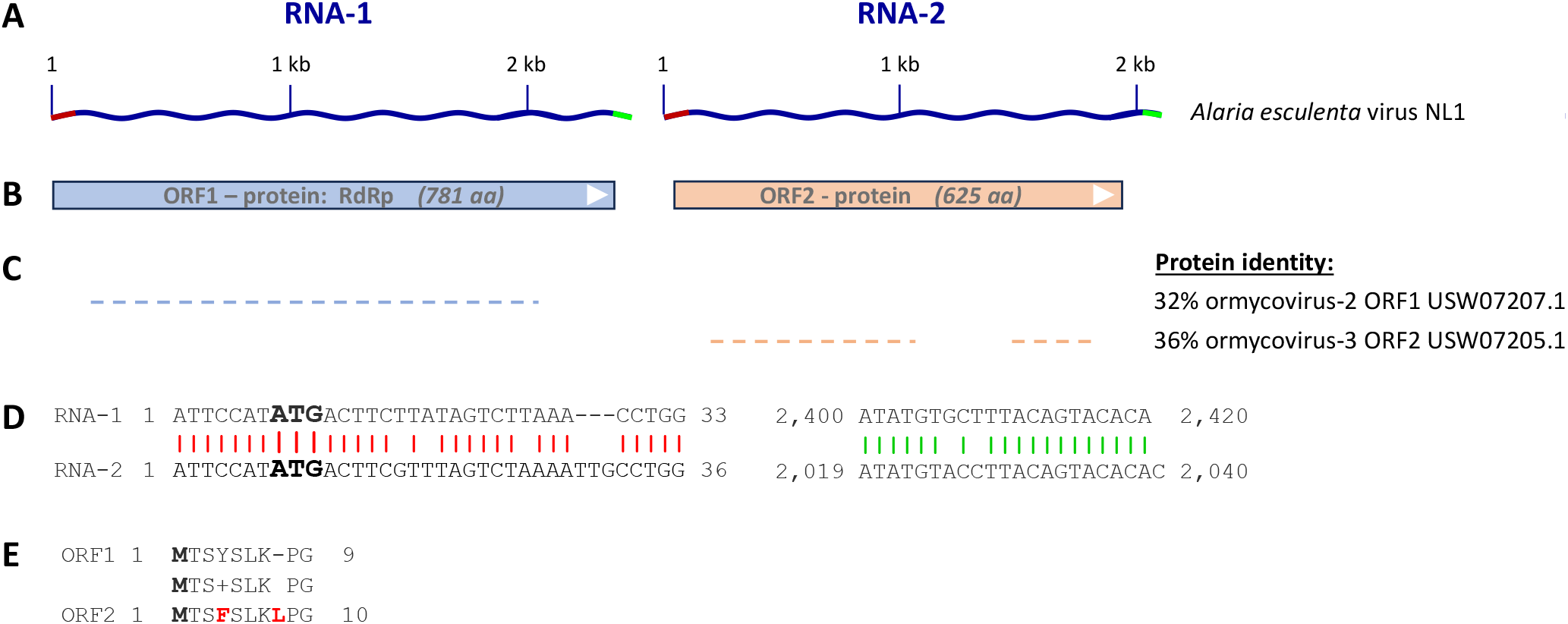
Characteristics of a new seaweed virus: Alaria esculenta virus NL-1 (Aev-NL1). A) Schematic representation of the Aev-NL1 bisegmented genome sequences RNA1 (PP793779) and RNA2 (PP793780); B) ORFs detected in the virus genome sequences; C) Similarity of the best BLASTP hits of each virus protein sequence in NCBI GenBank; D) Genome sequence alignment of the 5’ ends (∼33 nt; left) and 3’ ends (21 nt; right) of RNA1 and RNA2. E) Protein sequence alignment of the first 9-10 residues of ORF1 (RNA1) and ORF2 (RNA2) showing conservation of the N-terminal ends.

Noticeable with this new bisegmented virus was the sequence similarity at both ends between the two RNA sequences, which comprises ∼33 nt at the 5’ ends and 21 nt at the 3’ ends (Figure 2D) and is also found in some bisegmented ormycoviruses (Forgia *et al*., 2022). Since the translation start site is at position 8, the start codon for both ORFs lies in the identical 5’ sequence (Figure 2E).

Again, since all *A. esculenta* samples came from healthy individuals, there is likely no obvious disease phenotype associated with this new RNA virus.

A search of the NCBI-SRA repository with the Aev-NL1 RNA sequence yielded a similar, yet unknown bisegmented virus in an *S. latissima* RNA-seq experiment (NCBI-SRA: PRJNA564197; Li *et al*., 2020). Aligning the 5’ ends as well as the 3’ ends of the Aev-NL1 and Slv-NL1 RNA sequences supported the almost completeness of the new Slv-NL1 virus sequences (Supplementary file S3). The nearly completed new virus sequences (*Saccharina latissima* virus NL1, Slv-NL1; RNA1: PP793781, RNA2: PP793782) showed for both segments no sequence similarity to any known virus RNA sequence, but 87% similarity to the Aev-NL1 RNA sequences. There is weak protein similarity to known virus protein sequences Erysiphe lesion-associated ormycoviruses 2 (ORF1: 31%) and 3 (ORF2: 34%), but substantial similarity (91% and 83%) to the Aev-NL1 protein sequences (Supp. File S3).

### A new RNA virus in *Saccharina Latissima*: Slv-NL2

In all eight *S. latissima* samples from one location (South NL), we found sRNA evidence for a yet unknown virus (Table 1). Using bioinformatics analyses including assembly of the sRNA-seq reads into contigs, we were able to achieve for all but one sample (S12), a continuous sequence that appeared to be the full sequence of an approximately 4 kb unknown virus (Figure 3A; Supplementary File S4). A specific RT-qPCR confirmed the presence of viral RNA in all samples that were tested (n = 5, average Ct 16). The new virus variant genome sequences from the seven *S. latissima* samples were quite similar as they differed only two to nine nucleotides (Figure 3F). We arbitrarily named this new virus *Saccharina latissima* virus NL-2 (Slv-NL2; PP793783).

**Figure 3.**
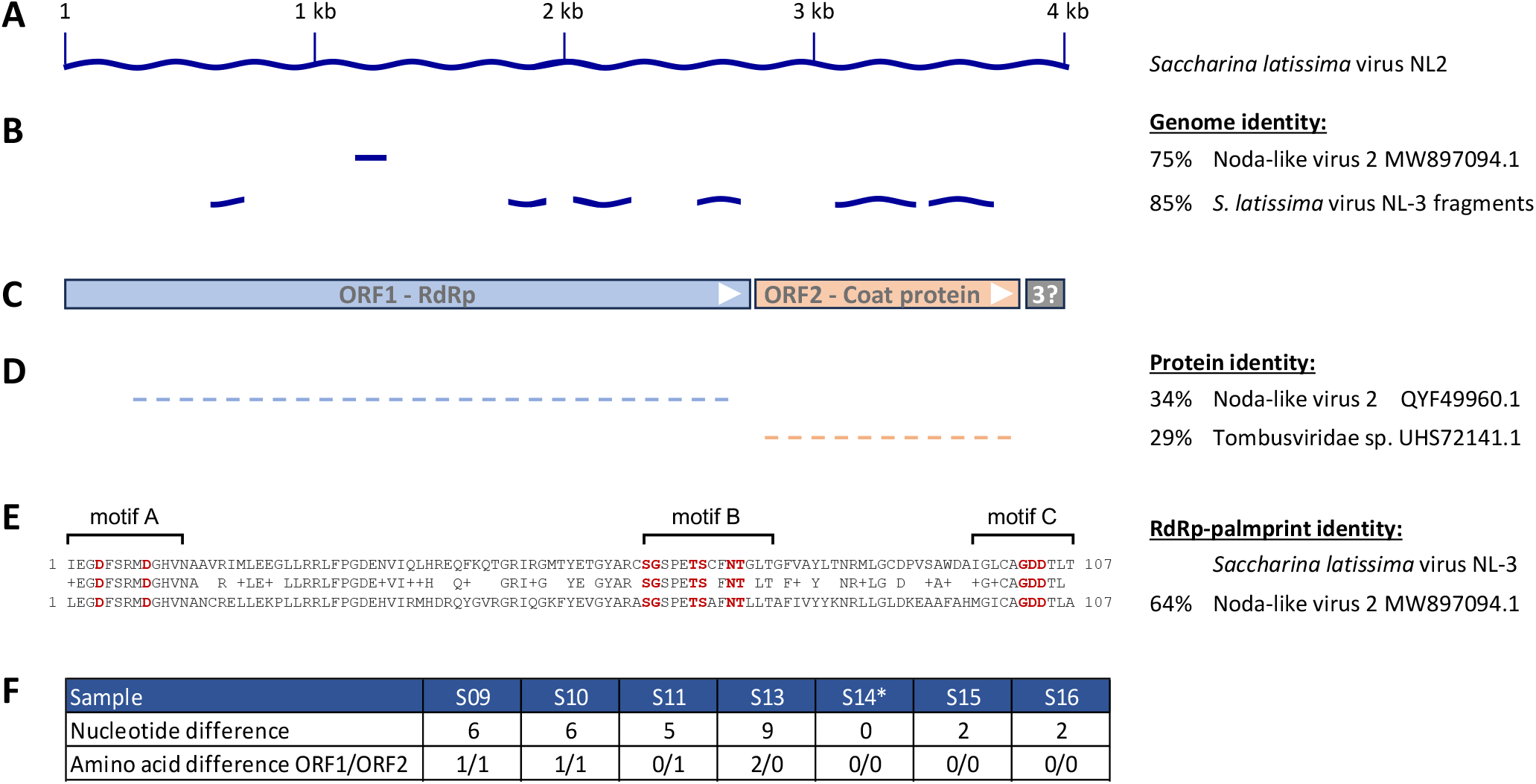
Characteristics of a new seaweed virus: Saccharina latissima virus NL-2. A) Schematics representation of the virus genome sequence (PP793783); B) Similarity of the best BLAST hit of the virus genome sequence in NCBI GenBank; C) ORFs detected in the virus genomic sequence; D) Similarity of the best BLASTP hit of the virus protein sequence in NCBI GenBank; E) RdRp palmprint comparison best match as determined with palmID webtool (serratus.oi, Babaian and Edgar 2022); F) Differences between the virus genomics sequences derived from samples S9-S13, S15, and S16 compared to S14 (*) plus the mapping sRNA-seq read counts for these samples, all others samples noteworthy read counts.

A BLAST similarity search on all virus genomic sequences in NCBI GenBank revealed almost no similar virus sequence. The longest similar sequence was a short stretch of 61 nucleotides from a noda-like virus (Figure 3B). Translation of the new SIv-NL2 RNA sequence revealed two clear ORFs of 909 aa and 349 aa and one potential ORF of 46 aa (without a stop codon; Figure 3C; Supplementary File S4). A similarity search revealed that the 5’ largest ORF (ORF1) appeared to code for the putative RNA dependent RNA polymerase (RdRp) gene, the 3’ smaller ORF (ORF2) appeared to code for the putative coat protein, and the utmost 3’ tiny putative ORF (ORF3) protein showed no similarity to any known virus protein (Figure 3C). The RdRp classification was substantiated by a high-confidence RdRp palmprint barcode score of 49.4 (Babaian *et al*., 2022) and a 64.2% identity match for the best matching RdRp in the database (Figure 3E).

As expected from the limited virus variant genome differences, the maximal differences between the SIv-NL2 variant protein sequences derived from distinct samples were just two and one amino acid, for ORF1 and ORF2, respectively, with half of the comparisons showing any difference (Figure 3F).

A BLASTP search on non-redundant protein sequences in the NCBI database revealed only 34% similarity with 88% coverage of SIv-NL2 ORF1 to a noda-like virus RdRp protein (isolated from river sediment in Jiangsu, China) and 29% similarity with 83% coverage of SIv-NL2 ORF2 to a tombus virus coat protein (isolated from river sediment in Heilongjiang, China) (Chen *et al*., 2022; Figure 3D; Supplementary File S4). Both virus families are non-enveloped positive-strand RNA viruses, with tombus viruses having a monopartite genome of about 4-5 kb, while noda viruses have a segmented genome of about 4 kb with the RdRp gene on RNA1 and the coat protein on RNA2. We did observe a potential third ORF at the 3’ end of the virus sequence that is missing a stop codon. It putatively codes for only a 45 aa protein, which is much smaller than the approximately 190 aa of the movement protein at that position in Tombusviridae.

Given the low similarity to known virus sequences, plus the fact that those known viruses are by themselves already only similar, such as “noda-like”, it is impossible to decisively classify this new virus into the virus taxonomy as it seems to unite tombus-like and noda-like viruses (Dolja & Koonin, 2018).

Additional bioinformatics analyses of the contigs assembled in this experiment using the Slv-NL2 sequences revealed six fragments (combined 1,427 nt) of a virus similar to Slv-NL2 (Figure 3B; Supplementary File S5), which we named *S. latissima* virus NL3. Slv-NL3 has a sequence identity of 85% compared to Slv-NL2 at a coverage of 35%. In contrast, the fragments similar to Slv-NL2-ORF1 and ORF2 showed an average protein similarity of 94% and 97%, respectively (Supplemental File S5). Just like for the Slv-NL1 virus, we discovered Slv-NL3 specific siRNAs in all eight samples from one sample location (Table 1). Also, a Slv-NL3 specific RT-qPCR showed ample presence of this virus in all tested samples (n=5, average Ct 21).

In contrast to what was found with Aev-NL1, the sRNA-seq reads associated with Slv-NL2, as well as Slv-NL3 showed no clear size bias (Figure 1A and 1B). If there is an siRNA response to these two viruses, it is hidden in the virus-RNA degradation products.

As for the co-presence of the alike Slv-NL2 and Slv-NL3 viruses, their high similarity at protein level plus their substantial difference at RNA level, suggest that they may function as each other’s decoy virus.

## Discussion

We discovered four previously unknown viruses in a small experiment (n = 24) with commercial seaweed species from two organizations in the Netherlands. The three RNA viruses: Aev-NL1 (PP793779 and PP793780), Slv-NL2 (PP793783), and Slv-NL3, plus one DNA virus; a Feldmannia-irregularis phaeovirus are described in detail. An additional new virus was discovered in a *S. latissima* NCBI-SRA dataset. We like to mention that even though the new virus sequences were constructed from sRNAs isolated from seaweed samples, it cannot be excluded that a discovered virus originates from non-seaweed virus hosts, such as fungi, that by itself also lived in or with the sampled algae.

The experiment focused on the possible involvement of viruses with the unhealthy phenotypes observed in seaweed plants. RNA virus Aev-NL1 and Feldmannia-irregularis phaeovirus only occurred in healthy *A. esculenta* plants. The similar RNA viruses Slv-NL2 and Slv-NL3 occurred in all *S. latissima* samples from one location irrespective of whether the plant showed bleaching or not. However, this does not preclude involvement of these viruses in the disease process, as it might well be that the plants without bleaching were already infected by the virus, but still at an asymptomatic stage. Possible pathogenic involvement is supported by the absence of virus RNA in *S. latissima* samples from the second location where no bleaching at all occurred, as well as the absence of a siRNA defense response as can be concluded from the lack of the siRNA hallmark 21-mer bias (Figure 1B) and the lack of the siRNA hallmark 5’ terminal T nucleotide (Table 2) on the virus sRNA’s of the virus-associated sRNAs. However, as our discovery bioinformatics heavily rely on similarity to known virus protein sequences (mainly RdRp), it will not detect any virus that had a divergent RdRp protein sequence (Charon *et al*., 2022), so we might miss out on viruses present in these samples. It might also well be that the phenotypes are derived from bacterial pathogens or symptoms from plants that are unhealthy due to non-pathogenic causes. A quick search between the sRNAs from plants with and without bleaching did not reveal any obvious pathogenic organism (results not shown). Altogether, a more extensive, better balanced, and better controlled experiment should be done to achieve more decisive results about the causes of the described unhealthy seaweed phenotypes.

## Supporting information

Supplemental File S3-Slv-NL1 sequences

Supplemental File S4-Slv-NL2 sequences

Supplemental File S5-Slv-NL3 sequences

Supplemental Table S1 Aev-NL1 similarities

Supplemental File S1-Feldmania sequences

Supplemental File S2-Aev-NL1 sequences

## Data availability

The raw sequence reads have been deposited in the NCBI Sequence Read Archive under BioProject accession number PRJNA1089059. The following genome sequences have been deposited in NCBI GenBank: Aev-NL1 RNA1 (PP793779), Aev-NL1 RNA2 (PP793780), Slv-NL1 RNA1 (PP793781), Slv-NL1 RNA2 (PP793782), and Slv-NL2 (PP793783).

## Supplemental information

Supplemental Figure S1: Images of seaweed samples with and without bleaching

Supplemental File S1: Partial new Feldmannia_irregularis-like virus DNA and protein sequences

Supplemental File S2: Alaria_esculenta virus-NL1 RNA and protein sequences

Supplemental File S3: Saccharina-latissima virus-NL1 RNA and protein sequences

Supplemental Table S1: Comparison of Aev-NL1 similar virus sequences

Supplemental File S4: Saccharina-latissima RNA virus-NL2 RNA and protein sequences

Supplemental File S5: Partial new S. latissima virus sequences

## Acknowledgments

This research was directly and indirectly funded by the Swammerdam Institute for Life Sciences of the University of Amsterdam.

