## Supplemental File S3-Slv-NL1 sequences for "Discovery of novel RNA viruses in commercially relevant seaweeds *Alaria esculenta* and *Saccharina latissima*"

### Supplemental File S3, *Saccharina latissima* (Slv-NL1) RNA virus-1 RNA and protein sequences

#### RNA1 (partial)

START codon

STOP codon

End sequence similarity

**Size	2,414 nt**

**Highest match	Alaria esculenta virus NL1 RNA1**

**(Aev-NL1-RNA1)**

**Percentage Coverage	98%**

**Percentage Identity	86%**

**>Saccharina_latissima-virus-NL1-RNA1**

CCGCGCAGACCGTTATGACTTCTTATAGTCTTAAACCAGGAGAGAAGGGTAACCTTTCTCTCCCCCAACCCTCAGGATTTCCTGACGGGTTGTTAGAAACTCTTCGTAGAACTCTACGTTCGAGTAGTGGTTCCTTGCTTGGCGGCAACCACTTTACACAAAATCTTCCGCGCAGACCGTGCGAAGGACATGTCGAGGAGCTTCGTGAGCAGCTCCCTGACTTTAGATACAGAAGCCTTAGAAAGTTCAAGGCTCTATGTGAAGAGAGGTGTAGCACTTGTGCTACCTCTTTAGTAACGTGCGCGGCATACCTCAAGGTGCATGCGCACATACACAAGTGTCCAAAATTTTGCATTCACTTGTTTAAAAACTTCACTCTTTATAAAGAGCAAAAGGTATATGATGCTTGCCAATCAATAAAATTTGCAGCATCCAGTTTAGCTCAGGGAGGTTATAATGACCTCGCTGAGAAGAAATTAGCAGAGATACCATTCGTAACTCAGCTAGAAGAAGAGGTTGTGAGCCATAATGCTTGGCCAAACCTCATTGCCTTCCCGCTCTGGAACTTGGAGCCTGGGATCGACAAAGAATCTAAGTTTGATCTTAGTTTCTTACCCCCCCCTACACTGAAGCAGTGTAATGGGTTTCGGAAGAAATTCCGTAGATATCTCGATAAGCATATGCCAGAGATAGTTGAAACTCTTTCGTACACAGAATGTATGAAAGTTGGCCCGAACAAATTTTATGATGACGGCGAAATTCGCAAAGATAGTGAGACTCCTATCAATGCCGATGGACCTTTTTTATATCAGTCCTTCATGACTGGGCCACTCTCCGTAAGGGAAGTGTGGCTCCCGACAAAAGCCTTTAAGGCCGCGTCGACGTGGTGGCATCGCGTTGCTGAGCAGCTGCTTCACAAGCGACCCCACTTAATACTCTCACAGGATCCTGAGGATGCTGCGAGAACTGTACGGAAAAGATTCAAACCGTGCAAATCTATAGATCTGAAGGGCTCAGGCCTTCAGTTCCCAATCGAGTACATCATTATAGTACTTGAACAACTAACTGAGCTCTTCCCAGAGATGGAGGAGCGCAAGGACATCGCAGTTGATTTACTGCAACGAATGTCCATATTTAAGGATAAGAAATTCCATATCCCAACTCGCGGAGTTGGACTCGGGTATTTTACAAATATCAAAGTCATGGTCATAGACTGTCTTCTCGAAGACTACTATGTTATTGCATCTTTCGACGACGATATGCTCGTCAAAGATACTCAGTACAACGCTTGTATTAAGCGACTGAAATCGTATGGCTTTCTCATTAACGAAGAAAAGTCAGGGCACCACTGGCCAATTAACCCGTGGTTCCTAAATGTAGGCATTCTCATGGACGAGGAGAGTGTCTTAGGCTACAGTACGTGTAATGCGTACATGGCAGCCTCATTTACTAAGCGCTATCATTGGGAGCGCAAAAGTATCCTTCAACAGGTTTACCCGGAGGATTCTCATTATATGGCCTTCCATTATGAGAAAATATTTGGATATGAATTCTTCAAAGGAGAATCCATTCAACACCCTAACAACGGCGGTTTATGTTGGTGGGCACGAGAGCTCGGTGGTAAAGACCAAGGAGCTTACCTTCAGGCGCATTTAATGCCCCAAAGGATATATGACGAGTGGAACGGGGACATCCCGTACCCCACTTTCGCTGCTGATACTATATCAGCTGCGGATAGAAAGCTACACCACTTTAAAAGGAAGCGGATGTACAAAGAGAAGACCATGATTTACTCATGGGATTTTTATCATCTCCACCCTCGCATGAGGTCTGGAGAATCCATAAAATCGCACTCAAGCGATCTTGATGGCATAACGCCTGTTTGGAGAGAAATATTACTCCTCAAGCATTTCGGGGTCTCCCATGGGAGCCTCGAACAAGACATACCCAAAATAGTGATACCACTATTGGCGGCAAAGTTCCCACTCTCGCGAGATCCGATCCAAGCTTTTGCTAGGAACGAGTATTGCGATTCAGAAAATGAACCACAAGGAATCGTCTGTGATGAAACACGTGAACGATTACATGCCATCTCTACTTCTAAGAGAATAGGCCACGAATACTTCTTCCAAAAGAGGGAAGAGGTGCAAAAGGACCCGGAGATACCTCCGGAGCCCCCTCCATGTCCCCAACTCGAAAATCTCTTGTTCGGGGATCTCGAATGGTCCGGTCCGGACCTGTCGAGTATTGAAATTCAGGCCCCCTATGAGGAATCGTCTGAATCTGAAGGAGAGGGTGACCTCTCCGACGTCGAGTCCGACATTATGTCGGACGTGATGGATTTCATCGACGATATTTAATTGAATGTCGTCGAAAAGCTATAATCCCTTTCTCAGGGTTTTGGCAGATATGTA

#### RNA2 (partial)

**Size	2,023 nt**

**Highest match	Alaria_esculenta-virus-NL1-RNA2 [Aev-NL1-RNA2]**

**Percentage Coverage	98%**

**Percentage Identity	86%**

**> Saccharina_latissima-virus-NL1-RNA2***

*atga*CTTCTTTTAGTCTTAAAACGCCTGGGGGAGAAAGACTCTCCCAGGCCATGGCCGAGGTAGCATGCTACCCGGCCAATGTACAAATCTCGCACTTAGAGCGCAAGATTACCGAAGAGCTGATTCTCCACTCTCCGGATGTAAACATAGCATCGTTTAAAGATGCTTGTAACTACTGCTCTCCCATTATTGAGGGAGCGGTATTGATTGAACCAGCTTTAAAAATGAAGCAGTTCGTGGACCGCGTTAGTGAATACTTCTATCGCGCTCAAACTTCGACAACTCTCAAGGATATTGAGGAGTTGCAATCCAGCATGCAACATCTGAGCATGACTGGAAATGCTAGTGTCCGAGATGATTTCTCGTTCACGATAGTGCCCGGCAAGCGTGGGGCCTACGAGATCCGTAAGTTAATACTAACTATACCGGATCTTTCAAAATCAGATCAGTCACATAGAACAAATCTGGTAGAACACATCAAGTTCGCTGCGACCTTGTTGAGTATCAATACTGAAGATATTAAAATTAATAGAATCAGTTCGTTGGGGACAACGTGTCCGTTACCCGCGAAATTAGGAGCTGCAATGACAGAAATGCTGACATTACACAGATCGTCCACTGCTTCTGCTGCAGTAGGACAGTTCAAACTGGAACGTGACCTTAGGGTCACGACCGGAGGTGCACTCGCCATTATGGCGCATTTGCACAAAAGAAACCCCTTTTACCGTAGAGACGGCAAAGGAAAATACGTCACGTCTGAGCTTCTCAAGACGGTCGTCAACAATGCATTTGGTCTCAATGAGTCCAGATGCTCACAGTTCTCGAAGAGTTTCTTCAAGGCTGTATTCAGAGCGATGGTCACGAATGACCTTGTTCGCATGCCTTCGAGCTTCAGTAAAAGCGCGAAGGTAACATTTGATGTCTCCTCACCTGAAGGCATCATGAGGAAGGCGGGTTATACACCGCTCATCCCAGACACGACCAAGATACTCTTGGTCACAACGACGGTCGATAAATTCGACCATCGACTCTTTGGGTCGTCTGGCGACAACAAAGACTATGAGGCTCCCGGGACTATGTCCGGGGATCCTATGGATCCAGTGGTGGAACCACCAGCACCTCCTGCCCCTAAGCAGAAGGTGACATTTGCTCCCGGAACATCAGAAGATGATAAAGGGAAGGGCAAGCCGTCTGATTGGACGGCTGTCAAGAAAAAGAATCCATCCAAAAAGGATCGGAGGAAAAAGTCTGTCGGGGATCTCCCGACAGGCCCTGCAGTAAGTGACAAGCCGAAGGGCGATGTCGCTAAGGCGCCAGTAAAACCGGCAGCCAAGGAAAAGGTTCTTCTTGAAACGATTTCCAAAATTGACTCTGCGCGAAACACGCACAAAGCCTTTGGGGCGGGGGTGAAACTCCTCCTCCCCTTCATCGACCCTAGCTCTAAGCAGTCGATGAAAGACCAACTCAAGTACAGTTACAAGAACTGTTCTGAGAAATCTCTACTCTTTTTCAAAGAGCAGAGAAACTTCGTTGCGAGCGCCGAAAAGACGTACGCAGTTCTTCAGGCTTCTAAGAACCCGAAGAGCAAAGCCAAAGCAGAGCACTACGTGCTAGCTAGGAACAGAATGTGCAACACTATGTTGGACATGAAATTTTCCGACCGTACAGGGACCTGTTACGAGTCATACTCTGATATTCCTTTGGGAATTCGGAGATATTTCGAAAAGGCTCTTTCTCGCCAAATGCGAGGCAAGCCTAAGAGCGATGACATCATTGAGCCATTGCTAAAGAAATCCAAGAGCGACACTGTCGCTATGGACGTAACGCCACCTACTCCGCCTGCGGAGTTGGAGGTAGAAGATTCCCCGTTGGACCTATAAGGTTAGAAGGGATTAGGTATTGAGTTGCTAACTCGTACCTTCAGTCGAGTCCCCTTAGGGGCTTGACTATTTATAGAACCTGACGCGTATGACGCGTGAGATTCGCAACTGCTCTCGGCATAAGAGAAAAGTGAATATGTAC

** First four nts inferred from identity with the 5’ ends of Aev-NL1 RNA1, Aev-NL1 RNA2 and Slv-NL1 RNA1.*

```
Aev-NL1-RNA1-p5                TTCCATATGACTTCTTATAGTCTTAAA---CCTGGA-GAGAGGGAAAACCTTTCTCTCCCCCAGCCCTCTGGCTTC
Slv-NL1-RNA1-p5        CCGCGCAGACCGTTATGACTTCTTATAGTCTTAAA---CCAGGA-GAGAAGGGTAACCTTTCTCTCCCCCAACCCTCAGGATTT
Aev-NL1-RNA2-p5               ATTCCATATGACTTCGTTTAGTCTAAAATTGCCTGGGGGGGAGAAGCTCTCCCAGGCGCTGGCCGAGGTAGCATGCT
Slv-NL1-RNA2-p5                          CTTCTTTTAGTCTTAAAACGCCTGGGGGAGAAAGACTCTCCCAGGCCATGGCCGAGGTAGCATGCT


Aev-NL1-RNA1-p3              CGACATCTAATTAGATGTTGTCGATAAGCTATAATCCCAGTT-AAGGGTTTTGGCAGATATGTGCTTTACAGTACACA
Slv-NL1-RNA1-p3              CGATATTTAATTGAATGTCGTCGAAAAGCTATAATCCCTTTCTCAGGGTTTTGGCAGATATGTA
Aev-NL1-RNA2-p3              ACCTTATGCGTATGACGTATAAGATTCTTGACTGCTCTCGGCATCAGAGAAAAGTGAATATGTACCTTACAGTACACAC
Slv-NL1-RNA2-p3              ACCTGACGCGTATGACGCGTGAGATTCGCAACTGCTCTCGGCATAAGAGAAAAGTGAATATGTAC
```

#### Protein ORF1

**Size	763 aa**

**Highest match	Alaria_esculenta-virus-NL1-ORF1 [Aev-NL1-ORF1]**

**Percentage Coverage	100%**

**Percentage Identity	91%**

**Best match GenBank	RdRp, Erysiphe lesion-associated ormycovirus 2 (USW07207.1)**

**Percentage Coverage	80%**

**Percentage Identity	31%**

**>Saccharina_latissima-virus-NL1-ORF1**

MTSYSLKPGEKGNLSLPQPSGFPDGLLETLRRTLRSSSGSLLGGNHFTQNLPRRPCEGHVEELREQLPDFRYRSLRKFKALCEERCSTCATSLVTCAAYLKVHAHIHKCPKFCIHLFKNFTLYKEQKVYDACQSIKFAASSLAQGGYNDLAEKKLAEIPFVTQLEEEVVSHNAWPNLIAFPLWNLEPGIDKESKFDLSFLPPPTLKQCNGFRKKFRRYLDKHMPEIVETLSYTECMKVGPNKFYDDGEIRKDSETPINADGPFLYQSFMTGPLSVREVWLPTKAFKAASTWWHRVAEQLLHKRPHLILSQDPEDAARTVRKRFKPCKSIDLKGSGLQFPIEYIIIVLEQLTELFPEMEERKDIAVDLLQRMSIFKDKKFHIPTRGVGLGYFTNIKVMVIDCLLEDYYVIASFDDDMLVKDTQYNACIKRLKSYGFLINEEKSGHHWPINPWFLNVGILMDEESVLGYSTCNAYMAASFTKRYHWERKSILQQVYPEDSHYMAFHYEKIFGYEFFKGESIQHPNNGGLCWWARELGGKDQGAYLQAHLMPQRIYDEWNGDIPYPTFAADTISAADRKLHHFKRKRMYKEKTMIYSWDFYHLHPRMRSGESIKSHSSDLDGITPVWREILLLKHFGVSHGSLEQDIPKIVIPLLAAKFPLSRDPIQAFARNEYCDSENEPQGIVCDETRERLHAISTSKRIGHEYFFQKREEVQKDPEIPPEPPPCPQLENLLFGDLEWSGPDLSSIEIQAPYEESSESEGEGDLSDVESDIMSDVMDFIDDI

#### Protein ORF2

**Size	619 aa**

**Highest match	Alaria_esculenta-virus-NL1-ORF2 [Aev-NL1-ORF2]**

**Percentage Coverage	98%**

**Percentage Identity	83%**

**Best match GenBank	Erysiphe lesion-associated ormycovirus 2 (USW07205.1)**

**Percentage Coverage	57%**

**Percentage Identity	34%**

**>Saccharina_latissima-virus-NL1-ORF2***

*mt*SFSLKTPGGERLSQAMAEVACYPANVQISHLERKITEELILHSPDVNIASFKDACNYCSPIIEGAVLIEPALKMKQFVDRVSEYFYRAQTSTTLKDIEELQSSMQHLSMTGNASVRDDFSFTIVPGKRGAYEIRKLILTIPDLSKSDQSHRTNLVEHIKFAATLLSINTEDIKINRISSLGTTCPLPAKLGAAMTEMLTLHRSSTASAAVGQFKLERDLRVTTGGALAIMAHLHKRNPFYRRDGKGKYVTSELLKTVVNNAFGLNESRCSQFSKSFFKAVFRAMVTNDLVRMPSSFSKSAKVTFDVSSPEGIMRKAGYTPLIPDTTKILLVTTTVDKFDHRLFGSSGDNKDYEAPGTMSGDPMDPVVEPPAPPAPKQKVTFAPGTSEDDKGKGKPSDWTAVKKKNPSKKDRRKKSVGDLPTGPAVSDKPKGDVAKAPVKPAAKEKVLLETISKIDSARNTHKAFGAGVKLLLPFIDPSSKQSMKDQLKYSYKNCSEKSLLFFKEQRNFVASAEKTYAVLQASKNPKSKAKAEHYVLARNRMCNTMLDMKFSDRTGTCYESYSDIPLGIRRYFEKALSRQMRGKPKSDDIIEPLLKKSKSDTVAMDVTPPTPPAELEVEDSPLDL

** First two aa inferred from identity between 5’ ends of Aev-NL-1-RNA1, Aev-NL-1-RNA2, Slv-NL-1-RNA1, and Slv-NL-1-RNA2*

#### Protein similarity ORF1

Query	= **Alaria_esculenta-virus-NL1-ORF1 [Aev-NL1-ORF1]**

Subject	= **Saccharina_latissima-virus-NL1-ORF1 [Slv-NL1-ORF1]**

| **Alignment statistics for match #1** | | | | | |
| --- | --- | --- | --- | --- | --- |
| Score | Expect | Method | Identities | Positives | Gaps |
| 1472 bits(3812) | 0.0 | Compositional matrix adjust. | 711/781(91%) | 752/781(96%) | 0/781(0%) |

```
Query  1    MTSYSLKPGERENLSLPQPSGFPDGLLNTLRRTLRSSSGSLLGGNHFVGNLPRRPCEGHV  60
            MTSYSLKPGE+ NLSLPQPSGFPDGLL TLRRTLRSSSGSLLGGNHF  NLPRRPCEGHV
Sbjct  1    MTSYSLKPGEKGNLSLPQPSGFPDGLLETLRRTLRSSSGSLLGGNHFTQNLPRRPCEGHV  60

Query  61   EELRKQLPDFRYRSLKKFKAVCEERCGVCATSLVSCAAYLKVHAHIHKCPKFCMHLFKNF  120
            EELR+QLPDFRYRSL+KFKA+CEERC  CATSLV+CAAYLKVHAHIHKCPKFC+HLFKNF
Sbjct  61   EELREQLPDFRYRSLRKFKALCEERCSTCATSLVTCAAYLKVHAHIHKCPKFCIHLFKNF  120

Query  121  TLYKEQKIYDACQSIKFAASSLAQGGFNNLAEKKLSEIPFVSQLEDEVVSHNAWPNLIAF  180
            TLYKEQK+YDACQSIKFAASSLAQGG+N+LAEKKL+EIPFV+QLE+EVVSHNAWPNLIAF
Sbjct  121  TLYKEQKVYDACQSIKFAASSLAQGGYNDLAEKKLAEIPFVTQLEEEVVSHNAWPNLIAF  180

Query  181  PLWNIEPGIDKESKFDLSFLPPPTQKQCNGFRKKFRRYLDKHMPEIVETLSYTECMKVGP  240
            PLWN+EPGIDKESKFDLSFLPPPT KQCNGFRKKFRRYLDKHMPEIVETLSYTECMKVGP
Sbjct  181  PLWNLEPGIDKESKFDLSFLPPPTLKQCNGFRKKFRRYLDKHMPEIVETLSYTECMKVGP  240

Query  241  NKFYDDGEIRKDSETAINSDGPFLYQSFMTGPLSVREVWLPTKAFKASSTWWHRVAEQLL  300
            NKFYDDGEIRKDSET IN+DGPFLYQSFMTGPLSVREVWLPTKAFKA+STWWHRVAEQLL
Sbjct  241  NKFYDDGEIRKDSETPINADGPFLYQSFMTGPLSVREVWLPTKAFKAASTWWHRVAEQLL  300

Query  301  HKRPHLILSQDPEAAARTVRKRFKPCKSIDLKGSGLQFPIEYIIIVLEQLAELFPEMEER  360
            HKRPHLILSQDPE AARTVRKRFKPCKSIDLKGSGLQFPIEYIIIVLEQL ELFPEMEER
Sbjct  301  HKRPHLILSQDPEDAARTVRKRFKPCKSIDLKGSGLQFPIEYIIIVLEQLTELFPEMEER  360

Query  361  KDIAVDLLQRMSIFKDKKFHIPTRGVGLGYFTNIKVMVIDCLLEDYYVIASFDDDMLVKD  420
            KDIAVDLLQRMSIFKDKKFHIPTRGVGLGYFTNIKVMVIDCLLEDYYVIASFDDDMLVKD
Sbjct  361  KDIAVDLLQRMSIFKDKKFHIPTRGVGLGYFTNIKVMVIDCLLEDYYVIASFDDDMLVKD  420

Query  421  NQYNACIKRLKSYGFLINEEKSGHHWPVNPWFLNVGILMDEETVLGYSTCNAYMAAAFTK  480
             QYNACIKRLKSYGFLINEEKSGHHWP+NPWFLNVGILMDEE+VLGYSTCNAYMAA+FTK
Sbjct  421  TQYNACIKRLKSYGFLINEEKSGHHWPINPWFLNVGILMDEESVLGYSTCNAYMAASFTK  480

Query  481  RYHWERKSILQQVYPEDSHYMAFHYEKIFGYEFFKGESIEHPHNGGLCWWSKELGGKDQG  540
            RYHWERKSILQQVYPEDSHYMAFHYEKIFGYEFFKGESI+HP+NGGLCWW++ELGGKDQG
Sbjct  481  RYHWERKSILQQVYPEDSHYMAFHYEKIFGYEFFKGESIQHPNNGGLCWWARELGGKDQG  540

Query  541  AFLQAHLMPQRIYDEWNGEIPYPTFAADTISSADRKQHHFKRKRMFKDKTMIYSWDYYHL  600
            A+LQAHLMPQRIYDEWNG+IPYPTFAADTIS+ADRK HHFKRKRM+K+KTMIYSWD+YHL
Sbjct  541  AYLQAHLMPQRIYDEWNGDIPYPTFAADTISAADRKLHHFKRKRMYKEKTMIYSWDFYHL  600

Query  601  HPRMRSGESIRSHSSDLDGITPAWREILLLKHFGVSHGSLEQDIPKAVIPLLADKFPLSR  660
            HPRMRSGESI+SHSSDLDGITP WREILLLKHFGVSHGSLEQDIPK VIPLLA KFPLSR
Sbjct  601  HPRMRSGESIKSHSSDLDGITPVWREILLLKHFGVSHGSLEQDIPKIVIPLLAAKFPLSR  660

Query  661  DPIQAFARSEFCDSENEPQGIVCDETRERLLAISRATKVGHEYFFQKREEVNQDLEIPPK  720
            DPIQAFAR+E+CDSENEPQGIVCDETRERL AIS + ++GHEYFFQKREEV +D EIPP+
Sbjct  661  DPIQAFARNEYCDSENEPQGIVCDETRERLHAISTSKRIGHEYFFQKREEVQKDPEIPPE  720

Query  721  PNPTPQLDNLLFGDLEWSGPDLSGIEIQAPYEESSESEGEEILSDAMSDCMSNISEFLDDI  781
            P P PQL+NLLFGDLEWSGPDLS IEIQAPYEESSESEGE  LSD  SD MS++ +F+DDI
Sbjct  721  PPPCPQLENLLFGDLEWSGPDLSSIEIQAPYEESSESEGEGDLSDVESDIMSDVMDFIDDI  781
```

#### Protein similarity-ORF2

Query	= **Alaria_esculenta-virus-NL1-ORF2 [Aev-NL1-ORF2]**

Subject	= **Saccharina_latissima-virus-NL1-ORF2 [Slv-NL1-ORF2]**

| **Alignment statistics for match #1** | | | | | |
| --- | --- | --- | --- | --- | --- |
| Score | Expect | Method | Identities | Positives | Gaps |
| 1068 bits(2763) | 0.0 | Compositional matrix adjust. | 522/626(83%) | 571/626(91%) | 1/626(0%) |

```
Query  1    MTSFSLKLPGGEKLSQALAEVACYPANVQISHIERKITEDLILFSSDENIASITDACNYC  60
            MTSFSLK PGGE+LSQA+AEVACYPANVQISH+ERKITE+LIL S D NIAS  DACNYC
Sbjct  1    MTSFSLKTPGGERLSQAMAEVACYPANVQISHLERKITEELILHSPDVNIASFKDACNYC  60

Query  61   TPHIEGSVIIEPALQMRQFVDRVSKFVYRAHTSTTLKEIEELQSSMQNLSMSGNTNEQED  120
            +P IEG+V+IEPAL+M+QFVDRVS++ YRA TSTTLK+IEELQSSMQ+LSM+GN + ++D
Sbjct  61   SPIIEGAVLIEPALKMKQFVDRVSEYFYRAQTSTTLKDIEELQSSMQHLSMTGNASVRDD  120

Query  121  FSFMIVPGKRGAYEIRKLILTIPDLSKSDQSHRTNLVEHIKFAATLLSINTEDIKVDRVS  180
            FSF IVPGKRGAYEIRKLILTIPDLSKSDQSHRTNLVEHIKFAATLLSINTEDIK++R+S
Sbjct  121  FSFTIVPGKRGAYEIRKLILTIPDLSKSDQSHRTNLVEHIKFAATLLSINTEDIKINRIS  180

Query  181  SLGTTCPLPAKLGGAMTEMLTLHRSSTASAAVGQFKLERDLRVTTGGALAILAHLHKRSP  240
            SLGTTCPLPAKLG AMTEMLTLHRSSTASAAVGQFKLERDLRVTTGGALAI+AHLHKR+P
Sbjct  181  SLGTTCPLPAKLGAAMTEMLTLHRSSTASAAVGQFKLERDLRVTTGGALAIMAHLHKRNP  240

Query  241  FYRRDGKGKYVTSELLKTVVNNAFGLNESRCSQFSKSFFKAVFRAIVTNDLVRVPSSFSK  300
            FYRRDGKGKYVTSELLKTVVNNAFGLNESRCSQFSKSFFKAVFRA+VTNDLVR+PSSFSK
Sbjct  241  FYRRDGKGKYVTSELLKTVVNNAFGLNESRCSQFSKSFFKAVFRAMVTNDLVRMPSSFSK  300

Query  301  SAKVTFDVSSPEGIMRKAGYTPLIPDTTKMLLVLTTVDKLDHRLFGSTGDHEEIEDTGQK  360
            SAKVTFDVSSPEGIMRKAGYTPLIPDTTK+LLV TTVDK DHRLFGS+GD+++ E  G
Sbjct  301  SAKVTFDVSSPEGIMRKAGYTPLIPDTTKILLVTTTVDKFDHRLFGSSGDNKDYEAPGTM  360

Query  361  PKDP-PATEVATSPPADTPKVTFAPGTSEDDKGKGKASDWTTVKKKNPSKKDRRKKSTGD  419
              DP        +PPA   KVTFAPGTSEDDKGKGK SDWT VKKKNPSKKDRRKKS GD
Sbjct  361  SGDPMDPVVEPPAPPAPKQKVTFAPGTSEDDKGKGKPSDWTAVKKKNPSKKDRRKKSVGD  420

Query  420  LPVGPKSNDKPQGYVMEAPAKPAAKKKVHLETISKLDSARNTHKAFGAGVKLLLPFIDPS  479
            LP GP  +DKP+G V +AP KPAAK+KV LETISK+DSARNTHKAFGAGVKLLLPFIDPS
Sbjct  421  LPTGPAVSDKPKGDVAKAPVKPAAKEKVLLETISKIDSARNTHKAFGAGVKLLLPFIDPS  480

Query  480  SKLSMKDQLKYSYKNCSEQSLLFFKEQRNFVASAEKTYAVLQASKNPKSKATAEHYVLAR  539
            SK SMKDQLKYSYKNCSE+SLLFFKEQRNFVASAEKTYAVLQASKNPKSKA AEHYVLAR
Sbjct  481  SKQSMKDQLKYSYKNCSEKSLLFFKEQRNFVASAEKTYAVLQASKNPKSKAKAEHYVLAR  540

Query  540  NRMCNSLLPMKFADRTGTCYKSYSEIPLGVRRYFEKALSREIQERPKSDDIIEPSLKKSR  599
            NRMCN++L MKF+DRTGTCY+SYS+IPLG+RRYFEKALSR+++ +PKSDDIIEP LKKS+
Sbjct  541  NRMCNTMLDMKFSDRTGTCYESYSDIPLGIRRYFEKALSRQMRGKPKSDDIIEPLLKKSK  600

Query  600  VDTVDMDVTPTTPDTEMEVPEPPLDL  625
             DTV MDVTP TP  E+EV + PLDL
Sbjct  601  SDTVAMDVTPPTPPAELEVEDSPLDL  626
```
