## Supplemental File S4-Slv-NL2 sequences for "Discovery of novel RNA viruses in commercially relevant seaweeds *Alaria esculenta* and *Saccharina latissima*"

### Supplemental File S4, *Saccharina latissima* RNA virus-2 (Slv-NL2) RNA and protein sequences

START codon

STOP codon

#### RNA

**Size	3,976 nt**

**Highest match	M****W897094.1 MAG: Jiangsu sediment noda-like virus 2 isolate 51-141_384017	hypothetical protein gene, partial cds**

**Percentage Coverage	2%**

**Percentage Identity	75%**

**>Saccharina_latissima-virus-NL2-RNA**

CACAAATAACTAATTTTGTCCGTGATAAAAGATGTATTTTAGCCTACAGGGGCTGATTAATGCGATACTAAGCCATATCTTTGTCATGTATGAACCAGATTTCAAACAGTATTACACTGTGTTCGGTTGCATTGCCGGACTCACCCTCATGTTCCTAGTTACTGGAATTGTGTGGGCTGTAGACCGTGTGAGAATTATGAAGGAATTGACTTTCAGAATCTCACGAACCCTAGAGAGAACCAAGATCGAGCCTGCTATGGCTGCTAACTCGGTCGCTTCTGTTTTCAATAGGATGGACGTTCCAACGTTGAATGTCAGCAGTAGACACACCCATGCGTATTGCGCAGCGGTGCGCACATCTGCAGTAACTTTCCTCCGCCAGTTCTGCCTTACTTTAGGCGTTCCAGAGTACTGTGTTTCACGTTCTCGTCGGGATGAAAATCTCGGGGTTAACGGTAGTAAGCTATGGTACATGGGAATAGACGTGGATAAGAAAGTTACGAATGATGTGCCAAAGGGAATTGCTACCCTAGTTGACACTGATTTTCACGATGACGACCTACCGGAGATGCTACTCGGGGCTCTCCATCCGATCGTCCTGTATACGATCAATCCACGAGTAGTGGCCAAGTCCAGCGGCGAGGTCACATACTACTTCAACACGAACAATGAGATAGTGATGCGCTATTCAGGTGGAGCTTTATGGCAACATAAACTCTGGGACTACACATCTGACATCATATGCGTGCGGAAAAGGGTGCTTGGCATCACTTTGAAGTATTCACTGTTTCGGGTGGAGAAGAAACAAATGGACGATCTTCGTACGATTGTCTGCTTGGCACCGCTGGGAACCTGGGTTGGGCTTTCGGCCATACTGGCAAGTATGCGTCTTAAGGCTAAGCCCCTTGAGAGACTGGAGCCCGTTCATAATGGATGGGCTCACCTCATCGACATCGGCACCACCGGTATAACTCACTCTATTGGGCGAGCTGGATCCTATCAGGAAGCCAGTTTTTCGGCCGTGGAACTTGCAGCGTGCAAGGGAGCTACCGGCATGAGCAACCATGATGCAATGGTATCAACCACTTGCCCATGGTTAGACGGGGATAGACTTCGCGGCTTAGTTGCTGCAGAATATTTCCGCGACGGTGACGTGAAACCGACGTATGTAACATCGGTTGAGGACAGTGTTCGCCATATCAGTGCTGCAGTTGAGACTGCAGGTGATGAGGAGGAACGACAAGGGGTAACCGGGTTTATGTCCCCCCTAGTTACAGGGGGTTGCTACTCACATAACACATCAACTTCGAACAACAAATGGGGAGTCAAGGCTAGGATCACCGACCTTAGTAATGATGATTCCACATTGCGTCCGTTTCCCACATCTTGTCTTAAGGAGTTTTTGGACTTCTTTTGCAAGAGTGGACCATTGACGCCCGTTGCGTATGATGAAGTGTATGCAAATCAGGTTCGCCCAACACAGCGTGCAATCCTGCGTGAGGGAGCAATGAAGGGTGCCGTGGAAGACACCACGGTGCGATCTTTCTTGAAGAAAGAATCCGTGCAGGGTATCAAGGACCCGAGGGTTATCTCAACCATCGCTCCACGATCGAAGCTAGAGTATTCTAAATACATGTTACCCGTTGGACGACTTATGAAGTCTTTCAGGTGGTATGGGTTTAAGACTCCGGTGGAGGTGGCCGCAGACGTGGCAGAGACGGTGGGTGGGTTGCCCTATGCGATAGAAGGGGATTTCAGCAGGATGGACGGGCATGTGAACGCAGCGGTGCGTATCATGCTTGAAGAGGGCCTGTTGCGAAGATTGTTCCCGGGGGATGAGAATGTTATTCAACTCCACCGGGAACAGTTTAAGCAGACTGGGAGGATACGAGGCATGACCTATGAGACGGGTTATGCCCGGTGTTCAGGCTCACCCGAGACCTCGTGTTTTAATACTGGGTTGACCGGATTTGTGGCATATTTAACTAATAGGATGTTGGGTTGCGACCCAGTATCCGCTTGGGACGCCATAGGTCTGTGTGCAGGTGACGACACGTTGACTCCTGGTCACATAGGGATTTCCCCCGAAACGCATGCTTTGACGTTTAGGAAAGCTGCCCGACATGTTGGACAAGTCTTGACTGGTGACATCCGAATTAACGGTGAACCAGTGAAGTTCTTATCTCGTGTGTTCGGGGGCGCATGGTACGGGAGTCCGAACAGTATGGCCGACCCCTTAAGGGTTATTGTCAAGGTGCACAGTACACCAAACATGCCTGCTGGGACGACACCCCAACAAAAGTGCTATGAGAAAGGTCTGTCTCTTTACCAGACCGATTCCAACACGCCTATTTTGGGCCCTTTGGCCCGGAAGATGATGAGCGTCGGGAAGCCAGGAAAAGTCACTTCGGATTTTGCACCCACCTCTTGGTGGTCACAATTCGAAAACTCATGGCCTAATCAGGCTGAGGACTGGATGAACGATGTCATTCATGAGCAACTTGAGGACTTCGACCTCGCGAAGTTCGAGTCGTGGATTGACGTTGGTGACGTTTACAACGCACCGACGTGCATGGCTCTCCCGGAGAAACCGGGTAGCGTGGATGTTTTAGTGGATGGTGAGCTGGTACGCGGCAAACCTAAGGAGAAAGATTGCAGTAATACTGTTTCAGACGTCAAGAAACGCGTACACAGGCGAAAATCTCGCAGTAGAACACGTTCAGTTAGTGGACGTAGTGATGTAAGTGCTAATACAGTGTAGTCCTGGTGGTAGAAATAAATATGGCACCGTCATTGAAGATTAAGAGGGGTAAAACAGTTAAACCTAAGCAGAAACTTCGTGTTTCTGGCAGTGGCTTGACTTCACTTGCGAGGCTCATTAATGATCCATGCAAGGCGCCGTTGGTGGCTCCACAATATGGAGCTTCGAGCGGTGGTTACCTAACCAAGCTTTCTAGCTTTTCGAAGATCGACTTTGCCAATGGGAGTGATGGGTGGCTTGTTTGGTTTCCTGATTATCATGGCCGCAAGGGCATTGCAGGTCAGGGGGCTAACATTTTCGTTTATCAGAACGATAATGCAGTGAACACGCCAAACAACACTTCAAGTAACCCTTTGGGAGGTTCCGACACTGCAGTCACGGGATTGGGTGGATGGTCTATCGACGATCCCGCCTATTCTTTTGTTGCCGGAACAATTGTGCAAGATGCCCGAGCGGCCGCTGCTTGTGTCAAACTTACCTATACTGGTAGGAATGATGCTTTGGCAGGTCGAGTCGGTGTGCTTGAAGGCGTACCTAGGGACGCTCTCCTTACTGGGGGTGGAGTGAATCCGCCTACAATTACTGAAATGTTTAATTATGCTGCTACGGTGTCGCGTACTCCTATGGATGAAGTTGAGCTAAAGTTCCGCCCAAGCGAAGGGTCCGAACTTTATCGCAATGCCACTCCATCTGATTCCGCTTTTGTAATCGGTGATAGTGCCCCGACTAGTATTGGTGAAGGAACCCCCACGGGAGTGGGGCAGGGTATCGGTTTCTGTTGGTCTGGAATGGACACGTCTACCTCCATAACTTTGGATTTTCTGAAGGTTTTGGAGTGGAGGCCTGAATTCCAGAGTGGCCTAGTGTCACCCCCCCAACATGTGGCTGATGGTGGCGGGAACATGGTTTCTCGCGCCATTGCCTATCTGGACCGTCATTTCCCTGGGTGGCAACACAATTTGGGGAAGATTGGTAGTTCCGTTGCGGCACGCATTGCGCAAACTGCATTTAAGGGACCGGCCAATGATCTTCTCCGTATTGGAGCTGCTGCTGCTCCTATGTTACTTTGATGTGATATGTCTGTATGGCTTAGTAGTCCTAAACTACTCTTCTTACGAAGTAAAATCATGAAAAGCGCAAGGCGCTCCGTTTATTCTCTACACAAACGGTTTTACTCCGGGGCAGTACCGGCCATACCTCTGGTTACGAGGACCA

#### Protein SLv-NL2-ORF1, putative RdRp

**Size	909 aa**

**Highest match	QYF49960.1 MAG: Jiangsu sediment noda-like virus 2 isolate 51-141_384017	hypothetical protein gene, partial cds**

**Percentage Coverage	88%**

**Percentage Identity	34%**

**>Saccharina_latissima-virus-NL2-ORF1**

MYFSLQGLINAILSHIFVMYEPDFKQYYTVFGCIAGLTLMFLVTGIVWAVDRVRIMKELTFRISRTLERTKIEPAMAANSVASVFNRMDVPTLNVSSRHTHAYCAAVRTSAVTFLRQFCLTLGVPEYCVSRSRRDENLGVNGSKLWYMGIDVDKKVTNDVPKGIATLVDTDFHDDDLPEMLLGALHPIVLYTINPRVVAKSSGEVTYYFNTNNEIVMRYSGGALWQHKLWDYTSDIICVRKRVLGITLKYSLFRVEKKQMDDLRTIVCLAPLGTWVGLSAILASMRLKAKPLERLEPVHNGWAHLIDIGTTGITHSIGRAGSYQEASFSAVELAACKGATGMSNHDAMVSTTCPWLDGDRLRGLVAAEYFRDGDVKPTYVTSVEDSVRHISAAVETAGDEEERQGVTGFMSPLVTGGCYSHNTSTSNNKWGVKARITDLSNDDSTLRPFPTSCLKEFLDFFCKSGPLTPVAYDEVYANQVRPTQRAILREGAMKGAVEDTTVRSFLKKESVQGIKDPRVISTIAPRSKLEYSKYMLPVGRLMKSFRWYGFKTPVEVAADVAETVGGLPYAIEGDFSRMDGHVNAAVRIMLEEGLLRRLFPGDENVIQLHREQFKQTGRIRGMTYETGYARCSGSPETSCFNTGLTGFVAYLTNRMLGCDPVSAWDAIGLCAGDDTLTPGHIGISPETHALTFRKAARHVGQVLTGDIRINGEPVKFLSRVFGGAWYGSPNSMADPLRVIVKVHSTPNMPAGTTPQQKCYEKGLSLYQTDSNTPILGPLARKMMSVGKPGKVTSDFAPTSWWSQFENSWPNQAEDWMNDVIHEQLEDFDLAKFESWIDVGDVYNAPTCMALPEKPGSVDVLVDGELVRGKPKEKDCSNTVSDVKKRVHRRKSRSRTRSVSGRSDVSANTV

#### Protein SLv-NL2-ORF2, putative coat protein

**Size	349 aa**

**Highest match	UHS72141.1 MAG: hypothetical protein 2 [Tombusviridae sp.]**

**Percentage Coverage	83%**

**Percentage Identity	29%**

**>Saccharina_latissima-virus-NL2-ORF2**

MAPSLKIKRGKTVKPKQKLRVSGSGLTSLARLINDPCKAPLVAPQYGASSGGYLTKLSSFSKIDFANGSDGWLVWFPDYHGRKGIAGQGANIFVYQNDNAVNTPNNTSSNPLGGSDTAVTGLGGWSIDDPAYSFVAGTIVQDARAAAACVKLTYTGRNDALAGRVGVLEGVPRDALLTGGGVNPPTITEMFNYAATVSRTPMDEVELKFRPSEGSELYRNATPSDSAFVIGDSAPTSIGEGTPTGVGQGIGFCWSGMDTSTSITLDFLKVLEWRPEFQSGLVSPPQHVADGGGNMVSRAIAYLDRHFPGWQHNLGKIGSSVAARIAQTAFKGPANDLLRIGAAAAPMLL

#### Protein SLv-NL2-ORF3

**Size	46 aa**

**Highest match	x**

**Percentage Coverage	x**

**Percentage Identity	x**

**>Saccharina_latissima-virus-NL2-ORF2**

MSVWLSSPKLLFLRSKIMKSARRSVYSLHKRFYSGAVPAIPLVTRT

#### Protein similarity

**Query	Protein SLv-NL2-ORF1**

**Subject	QYF49960.1**

| **Alignment statistics for match #1** | | | | | |
| --- | --- | --- | --- | --- | --- |
| Score | Expect | Method | Identities | Positives | Gaps |
| 387 bits(994) | 8e-125 | Compositional matrix adjust. | 283/839(34%) | 417/839(49%) | 49/839(5%) |

```
Query  83   SVFNRMDVPTLNVSSRHTHAYCAAVRTSAVTFLRQFCLTLGVPEYCVSRSRRDENLGVNG  142
            SVF   ++  L V   HTH   AA R+SA  F+     +LG     +  S  D+  G
Sbjct  34   SVFAEAELSRLPVVGGHTHGVSAAARSSASAFIDTLAPSLGKRVVYIQGSSADQRKGRVY  93

Query  143  SKLWYMGIDVD------KKVTNDVPKGIATLVDTDFHDDDLPEMLLGALHPIVLYTINPR  196
            ++ +  G D++      +K  ND    +  ++D D+H D +P+ L     P++LYT+ P
Sbjct  94   TRHYRWGKDLNVTPRQVEKSEND----MTAMIDVDYHVD-MPKHLARNFKPLILYTLQPA  148

Query  197  VVAKSSGEVTYYFNTNNEIVMRYSGGALWQHKLWDYTSDIICVRKRVLGITLKYSLFRVE  256
                S+GE  Y F+    +    SGG  +QHKLW++  D +   + V  I L YS+F +E
Sbjct  149  RAGSSTGEYKYCFDAEGNVKYFVSGGGEYQHKLWNWKGDSVSATRSVCCIPLTYSVFAIE  208

Query  257  KKQMDDLRTIVCLAPLGTWVGLSAILASMRLKAKPLERLEPVHNGWAHLIDIGTTGITHS  316
            +KQ+D+   IV +APL  + G+   +A +R KA  L+RL+PV   +  ++  G +G+T S
Sbjct  209  RKQVDEDHQIVLVAPLSKYKGVYCWIAKLRAKAPELKRLDPVDGLFVRVLANGPSGMTVS  268

Query  317  IGRAGSYQEASFSA---VELAACKGATGMSNHDAMVSTTCPWLDGDRLRGLVAAEYFRDG  373
              R G +   +        +A+    T    H  + +        D     V  EY   G
Sbjct  269  TSRVGGFLSCNVPVNVDEAIASAANTTAKITHATLKAKMAQATSDDFTGSEVLLEYHLRG  328

Query  374  DVKPTYVTSVEDSVRHISAAVETAGDEEERQGVTGFMSPLVTGGCYSHNTSTSNNKWGVK  433
              K   +  V D+VR           E E+  +  FMSP V G     N   +N+K  V
Sbjct  329  CPKVERIDIV-DAVRSFQWVKTYQEYEPEKPAMVAFMSPFVHGAFVPDNC-LNNDKRMVS  386

Query  434  ARITDLSNDDSTLRPFPTSCLKEFLDFFCKS-GPLTPVAYDEVYANQVRPTQRAILREGA  492
             RI  L     TL PF  +C+K+F+  F +  G L PV  +EVYA Q +P+QR IL E A
Sbjct  387  ERIEKLKKPAQTLTPFLDACIKDFVKQFKEEVGMLWPVDNEEVYARQSKPSQRHILDE-A  445

Query  493  MKGAVEDTTVRSFLKKESVQGIKDPRVISTIAPRSKLEYSKYMLPVGRLMKSFRWYGF-K  551
              G     T  SF K E+   + DPRVISTI    K+EYS ++  +   +K+F WY F K
Sbjct  446  QHGRPNGKTA-SFEKNEAYPSVNDPRVISTINGVDKMEYSAFIYALADALKNFGWYAFGK  504

Query  552  TPVEVAADVAETVG-GLPYAIEGDFSRMDGHVNAAVRIMLEEGLLRRLFPGDEN--VIQL  608
             P E+A  VA+     L +    DFSRMDG VN   R  LE  L+  LF    +  +I+L
Sbjct  505  KPRELAERVAQICELALSHVDLTDFSRMDGRVNELAR-YLERLLMLALFSVKHHLALIKL  563

Query  609  HREQFKQTGRIR-GMTYETGYARCSGSPETSCFNTGLTGFVAYLTNRMLGCD-----PVS  662
               Q    GR + G+TY++GYAR SGSPETS FNT L  F+AYL  RM   D
Sbjct  564  MNSQTGLRGRTKHGVTYDSGYARASGSPETSAFNTILNAFIAYLAFRMTRRDGRYMTHKE  623

Query  663  AWDAIGLCAGDDTLTPGHIGISPETHALTFRKAARHVGQVLTGDIRINGEP-VKFLSRVF  721
            +W ++G+  GDD +T    G + E       KAA  +GQ LT D  I G P V FL+R +
Sbjct  624  SWTSLGVYGGDDGMTADVDGKAAE-------KAASMMGQKLTSDRVIRGFPGVTFLARHY  676

Query  722  G-GAWYGSPNSMADPLRVIVKVHSTPNMPAGTTPQQKCYEKGLSLYQTDSNTPILGPLAR  780
            G   W GSP S  D  R + K H T  + +  TP+ K  EK  + + TD+NTPI+G   +
Sbjct  677  GPDVWLGSPISCCDISRQLSKFHVTIRLSSKITPEIKLKEKSFAFHLTDANTPIIGEFVQ  736

Query  781  KMMSVG--KPGKVTSDFAPTSWWSQFE--NSWPNQAEDWMNDVIHEQLEDFDLAKFESWI  836
            +++++   KP +  ++     W  + +  N +PN+ EDWM D++  QL D+++  F  W+
Sbjct  737  RVLTLYPLKPRQFKNNL--NIWAVELDSSNQYPNEYEDWMLDLVKLQLPDYNIDSFRDWL  794

Query  837  DVGD---VYNAPTCMALP-EKPGSVDVLVDGELVRGKPKEKDCSNTVSDVKKRVHRRKS  891
               D   ++  P   A P   P    V VDG+ V+ + K    +   +  +K+ HR ++
Sbjct  795  LTADHATIFEMPNFGAQPYPNPKEGVVAVDGDFVKVEEKTPKITEARATPRKKFHRARN  853
```

#### Protein similarity

**Query	Protein SLv-NL2-ORF2**

**Subject	UHS72141.1**

| **Alignment statistics for match #1** | | | | | |
| --- | --- | --- | --- | --- | --- |
| Score | Expect | Method | Identities | Positives | Gaps |
| 122 bits(306) | 4e-36 | Compositional matrix adjust. | 87/301(29%) | 147/301(48%) | 17/301(5%) |

```
Query  30   ARLINDPCKAPLVAPQYGASSGGYLTKLS-SFSKIDFANGSDGWLVWFPDYHGRKGIAGQ  88
            A ++ +PC A L    +G S  G+L +   S+     A  + G+++W P+YH      G
Sbjct  46   ALMVANPCSAQLHPGLFG-SEEGFLGRFKVSYPLPSAAGATCGYVLWAPNYHNGGVNGGV  104

Query  89   GANI---FVYQNDNAVNTPNNTSS-NPLGGSDTAVTGLGGWSIDDPAYSFVAGTIVQDAR  144
                   F++Q+++  + P NT++ N   G+    + +   +I DPAY FV G   +DAR
Sbjct  105  AGGTGNVFIFQSNSPASAPTNTTAGNSAFGAGLTTSTVTASTIADPAYGFVNGATCRDAR  164

Query  145  AAAACVKLTYTGRNDALAGRVGVLEGVPRDALLTGGGVNPPTITEMFNYAATVSRTPMDE  204
              +AC+++ Y G   +++G+V ++E +P   LL     N P++  +F+Y+    R  +D
Sbjct  165  TVSACIRMDYLGAISSVSGQVAMVENLPMTDLLQ----NLPSVNNLFDYSTKSQRVSLDT  220

Query  205  VELKFRPSEGSELYRNATPSDSAFVIGDSAPTSIGEGTPTGVGQGI-GFCWSGMDTS--T  261
             E+ FR ++   L R    +D A ++G +   S        +   + GF + G+
Sbjct  221  SEIIFRSTQ-LGLDRFLAEADQALLVGATGAVSTVAPDAERIEPVVFGFAFRGVTAGELA  279

Query  262  SITLDFLKVLEWRPEFQSGLVSPPQHVADGGGNMVSRAIAYLDRHFPGWQH-NLGKIGSS  320
                +F+K +EWRP+   GL +        G NM S AIA LD H PGW      K+ S+
Sbjct  280  KFNFEFVKNMEWRPQTGQGLSATTS--TSTGVNMHSTAIAVLDHHAPGWTTPTYNKVASA  337

Query  321  V  321
            V
Sbjct  338  V  338
```
