## Supplemental File S5-Slv-NL3 sequences for "Discovery of novel RNA viruses in commercially relevant seaweeds *Alaria esculenta* and *Saccharina latissima*"

### Supplemental File S5, Partial *Saccharina latissima* RNA virus-3 (Slv-NL3) RNA and protein sequences

#### RNA fragments

**Size	906 nt**

**Highest match	Slv-NL3**

**Percentage Coverage	24%**

**Percentage Identity	85%**

**>Saccharina_latissima-virus-NL3-RNA**

>Fragment-1

ATCATCAGCTGAGACTTCTCACACGTTCAATGACGACGGCAAGATTTCCGTCGACATCGCCGGGGGAGCT
CACTACGAGCATTACCTTTGGGACTGGACCCCGGACATCATTACGTGTTCAAAGAGAATCTTCGGCGTCC
CCTACGCTACGACTGTCTACAAAGTCGAGAAACGCACTATCAACGAGCTCAAGGCACTTGTCTTGCTCGC
GCCCACTGGAACATGGTACGGACTCAACGCGATCGTGGCCAGCTTGCTGCAATCTTCTGCACTCAGA

>Fragment-2

GCATGTGAACGCAGCGGTGCGGACAATGCTCGAAGAGGGCCTGTTGCGGAGGTTGTTCCCAGGGGATGAGAATGTAATTCAACTCCACCGTGAACAATTTAAGCAAACTGGGAAGATACGTGGTATGACCTATGAGACGGGTTAT

>Fragment-3

TGATACTCATGGGCTTGAGAATTGTTGAGGCAGGCGAATGTCGCGAATGGGCGGCTGTGAGCCAGCGGAA
GCATTCGACGCAATTGGCATTGTGGCGGGTGACGACTCACTGAAGCCGGGCCACCGGAACGTAGAAGCCT
CCGTAGGCGGCAAATTCTTCGAACGCGCTGCTCGTCAGATGGGACAGAAGCTCACATCGGACATCAAGCA
CATCGGACAACCTGTACAATTTTTGGCCAGGGTG

>Fragment-4

GGACTTCGACCTCGCGAAGTTCGAGTCATGGATCACTGACGGTGACGTTTACAACGCCCCAACGTGCATGACTCTCCCGGATAAACCGGGTAGCGTGGATGTTTTAGTGGATGGTGAGCTGGTACGCGGCAAACCTAAGGAGAAAGATTGCAGTAATAC

>Fragment-5

GACAATGCAGTGAACACGCCTAACAACACTGCAAGTAACCCGTTGGGAGGTTCCGATACCGCAGTCGAGGGGCTTGGAGGGTGGTCTATTGATGACCCTGCTTACTCTTTCGTGGCCGGAACTATTGTGCAAGATGCACGTGTAGCCGCTGCTTGTGTGAAACTTACCTATACTGGTAGGAATGATGCTTTGGCTGGCCGTGTGGGTGTGCTCGAAGGTGTCCCTAGGGATGCTCTCCTGACTGGGGGCGGAGTCAATCCGCCTACCATTACTGAAATGTTTAACTATGCTGCCACGGTGTCGCGAACTCCTATGGATGAAGTTGAGCTAAAGTTCCGCCCAAGCGAAGGGTCCGAACTTTATCGCAATGCCACTCCATCTGATTCCGCTTTTGTAATCGGTGATAGTGCTCCGACGAGTATTGGTGAAGGAACCCC

>Fragment-6

CTGGAATGGACACGTCTACCTCCATAACTTTGGATTTTCTGAAGGTTTTGGAGTGGAGGCCTGAGTTCCAGAGTGGTCTTGTGGCCCCTCCCCAACATGTGGCTGACAGCGGTGGGAACATGGTTTCACGCGCTATTGCTTACTTGGACCGTCATTTCCCTGGAT

#### Protein Fragment-1, putative RdRp

**Size	92 aa**

**Highest match	Slv-NL2-ORF1**

**Percentage Coverage	9%**

**Percentage Identity	44%**

**>Saccharina_latissima-virus-NL2-Fragment1**

SSAETSHTFNDDGKISVDIAGGAHYEHYLWDWTPDIITCSKRIFGVPYATTVYKVEKRTINELKALVLLAPTGTWYGLNAIVASLLQSSALR

#### Protein Fragment-2, putative RdRp

**Size	48 aa**

**Highest match	Slv-NL2-ORF1**

**Percentage Coverage	5%**

**Percentage Identity	96%**

**>Saccharina_latissima-virus-NL2-Fragment2**

HVNAAVRTMLEEGLLRRLFPGDENVIQLHREQFKQTGKIRGMTYETGY

#### Protein Fragment-3, putative RdRp

**Size	80 aa**

**Highest match	Slv-NL2-ORF2**

**Percentage Coverage	7%**

**Percentage Identity	51%**

**>Saccharina_latissima-virus-NL2-Fragment3**

DTHGLENC-GRRMSRMGGCEPAEAFDAIGIVAGDDSLKPGHRNVEASVGGKFFERAARQMGQKLTSDIKHIGQPVQFLARV

#### Protein Fragment-4, putative RdRp

**Size	52 aa**

**Highest match	Slv-NL2-ORF2**

**Percentage Coverage	5%**

**Percentage Identity	92%**

**>Saccharina_latissima-virus-NL2-Fragment4**

DFDLAKFESWITDGDVYNAPTCMTLPDKPGSVDVLVDGELVRGKPKEKDCSN

#### Protein Fragment-5, putative coat protein

**Size	145 aa**

**Highest match	Slv-NL2-ORF2**

**Percentage Coverage	41%**

**Percentage Identity	98%**

**>Saccharina_latissima-virus-NL2-Fragment3**

DNAVNTPNNTASNPLGGSDTAVEGLGGWSIDDPAYSFVAGTIVQDARVAAACVKLTYTGRNDALAGRVGVLEGVPRDALLTGGGVNPPTITEMFNYAATVSRTPMDEVELKFRPSEGSELYRNATPSDSAFVIGDSAPTSIGEGT

#### Protein Fragment-6, putative coat protein

**Size	54 aa**

**Highest match	Slv-NL2-ORF2**

**Percentage Coverage	15%**

**Percentage Identity	96%**

**>Saccharina_latissima-virus-NL2-Fragment4**

GMDTSTSITLDFLKVLEWRPEFQSGLVAPPQHVADSGGNMVSRAIAYLDRHFPG
