## Supplemental Figure S1 for "Discovery of novel RNA viruses in commercially relevant seaweeds *Alaria esculenta* and *Saccharina latissima*"

### Slide 1
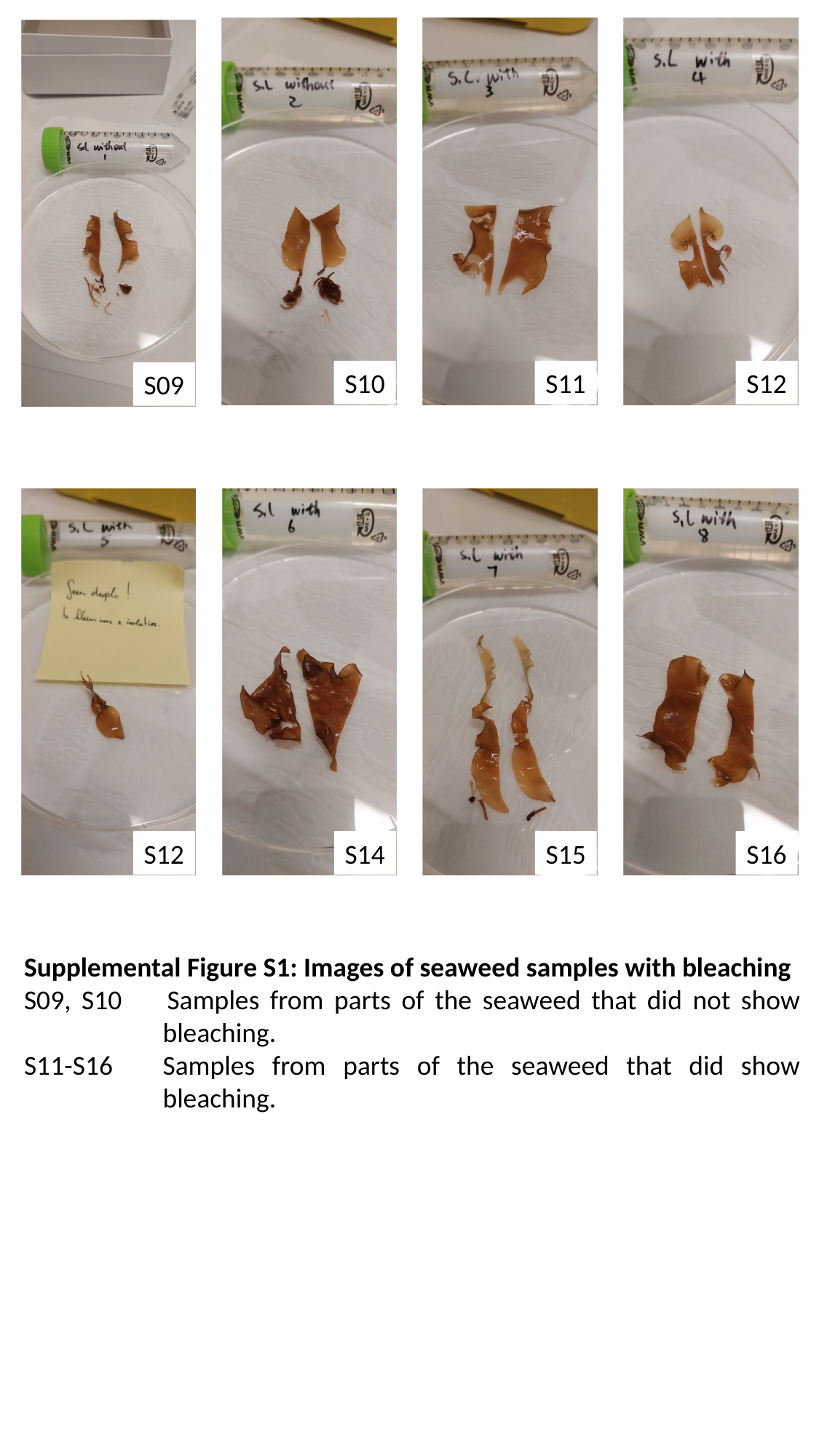

S10
S11
S12
S09
S12
S14
S15
S16
Supplemental Figure S1: Images of seaweed samples with bleaching
S09, S10	Samples from parts of the seaweed that did not show bleaching.
S11-S16	Samples from parts of the seaweed that did show bleaching.
