## Supplemental File S1-Feldmania sequences for "Discovery of novel RNA viruses in commercially relevant seaweeds *Alaria esculenta* and *Saccharina latissima*"

### Supplemental File S1, Partial, putative new *Feldmannia irregularis*-like virus DNA and protein sequences

#### DNA (partial)

**Highest match	NC_011183.1, Feldmannia species virus, complete genome**

**Percentage Coverage	66%**

**Percentage Identity	67%**

**>Two_siRNA_contigs_with_similarity_to_****Feldmannia_irregularis_virus_ major_capsid_protein**

AACAAACATGCCTGCCGGAGGAGGAGTCATCCAGATTGCCGCTGTCGGACGACAGAACGCCCACCTTAATGGAAACCCCGACCACACTCTTTTCAAGACCCAGCACCGTCGTTACACTTCGTTCGCTGAGGATCTTGAATACAATGACTTCAGCTCGGGAACCGTCGGATTCGGACAGAAGGTGTCTGCTTCCATCAGCCGTTACGGTGACCTTGTCTCGGACATCATGCTTGAAGTGAGCCTACCCGCCATCGAAGCCCCCCTTGAGGCCGCTTACATCACC

nn

TATCATTCCGGCTCGTACAGGCCTCACTGCCACCGACGCATCCGGTGTAGCACTTGCCGCCGGTGTCACCACTGGTTTCTCGGCTTACTGGGTGAACGCCATCGGTTTCGCTCTTATCTCTGAAGTACAGATTGAAATCGGAGGTACCGAAATCGACACTCTTTACCCCGAATGGATGTACTTCTGGGAGGAAATGACCCAGCGCCCCGGTGCCCGCCTCGGTGAACAGATCGGTAAGTTCTCGTACTCCGACACCGTCGAAGCTGACATGGTTGAATTCGCCAGCCAGGACCGCACTCTTTTCGTGCCCCTCCAGTTCTGGTTCAACAAGTACTTCATGGAACACGGTCTTTCGATCCCTCTGATCGCTCTTTCGTACCACGAAATCCGTGTACGCGTGACTTTCCGTTCCATCGCTGAATGCACCGTGTGTACTTTCACCGACGCCTCTCTTAACGGGGGTGGTGGTGTCAACGACATCGAACAGGACATCGAGACCCTTGTGTCCGGACTTGTACCCGTCAACAGCAACACCGCTTCCACCCTTGTAAGCGCCGACCTTGAAGCCCGTCTTCTTGTTTCGTACGTGTACCTTGACGCCGAAGAGCGTAACGCTTTCTCGTCCGTTGAACACGAATACCTCATCACCACCGTACAGCGCGCCCAGCACAACATCACCAGTGCTAACGCCACCTCCGACCAGGTCAAGATCTTCTTCAACCACCCCAGCAACTGCCTCACCTGGATGATCCGTCCTTCCGAGTACCTTACCGGTGCCGCCCGCCGCCGTTACTCCGTCGGACACAAGGACATGTTCGACTTCTCGCTCGCCAGCGGAAGCGCCGCAGTGGGACTTCCCTTCGGTGATGTGACCGACGCCACCAAATCGGCTTCGCTCACCCTTAACGGACACAGCCGCTGGCCCACTGACCTACCCGGTCTGTTCTTCCGTCAGACTGTGCCCGTCATGAAGTGGACCAACGCTTCCGACGGTTACATGTACGTGTTCAACTTCTCCGCCCGCGGTGGAGCATGGCAGCCAACCTCCACCCTTAACATGAGCCGTATCGACCACGTACAGCTTGAACTCAAATACGGCGCATCCATGCCCGTGTCTGACGTACTCATCTTCGCTGAGTCGTATAATCTGTTCATCGTTAAAGAAGGGATGGGCGGTATCAGATACTCCAACTAAGACCGTTCTTGTGGGGGTATGGTTTAATACTATTCCCCTACTATATATGTTTAAAAAATCTATCGTAAAATATGATCTAATTCAAAGAAATCATATTGAGACTTTAGTGGCAACCTCGACCTGTGCTGTATATTAAATTAAAATTAACAAAATATTTCAAACAAAAAAAAATTT

#### Protein (partial)

**Highest match	YP_009665663.1, FirrV-1-B50 [Feldmannia irregularis virus a]**

**Percentage Coverage	94%**

**Percentage Identity	66%**

**> Protein_with_similarity_to_Feldmannia_irregularis_virus_major_capsid_protein**

MPAGGGVIQIAAVGRQNAHLNGNPDHTLFKTQHRRYTSFAEDLEYNDFSSGTVGFGQKVSASISRYGDLVSDIMLEVSLPAIEAPLEAAYIT

X

IIPARTGLTATDASGVALAAGVTTGFSAYWVNAIGFALISEVQIEIGGTEIDTLYPEWMYFWEEMTQRPGARLGEQIGKFSYSDTVEADMVEFASQDRTLFVPLQFWFNKYFMEHGLSIPLIALSYHEIRVRVTFRSIAECTVCTFTDASLNGGGGVNDIEQDIETLVSGLVPVNSNTASTLVSADLEARLLVSYVYLDAEERNAFSSVEHEYLITTVQRAQHNITSANATSDQVKIFFNHPSNCLTWMIRPSEYLTGAARRRYSVGHKDMFDFSLASGSAAVGLPFGDVTDATKSASLTLNGHSRWPTDLPGLFFRQTVPVMKWTNASDGYMYVFNFSARGGAWQPTSTLNMSRIDHVQLELKYGASMPVSDVLIFAESYNLFIVKEGMGGIRYSN

#### **FirrV-1-B50 [Feldmannia irregularis virus a**]

**Sequence ID:**	[**YP_009665663.1**](https://www.ncbi.nlm.nih.gov/protein/YP_009665663.1?report=genbank&log$=protalign&blast_rank=1&RID=WTNK24MJ114)

**Size: 	435 aa**

>YP_009665663.1 FirrV-1-B50 [Feldmannia irregularis virus a]

MTLFKTVHRRFTSFAEDLEENDFSAGTVGFGQKVSATISRYGDLVTDMFMEIHLPPIEAAVNVTDASGNTVADADKAAYWVNAIGFALISEVQIEIGGTEVDILYPEWMFFWEEMTQRPGARLGEQIGKFTYSADVEEDMIEFAQQARTLYVPLPFWFNKYFMEAGLSIPLIALTYHEIKVKVTFRPLSECCCVVYRTEDPTHGEYYALAEGKVPVNSTSGSTLVSSDMDAKLLVSYVYLDKKERDAFATTEHTYLITTTQRQMHAITSAGSSSDQIKLYFNHPSNCLTWFVRPTDWMTNRRRYSVGHMDLFDYSLKSTSDVSVWGDVTDPVKTASLTLNGHSRFPDNMPGLFFRQTQPIMKWPNCSNGYMYVFSFSLQGGSWQPTSTLNMSRIDHVQLELKYGTSIPTSNVFVFAESYNLLVVKEGMGGVRYSN

#### Protein similarity

| **Alignment statistics for match #1** | | | | | |
| --- | --- | --- | --- | --- | --- |
| Score | Expect | Method | Identities | Positives | Gaps |
| 635 bits(1637) | 0.0 | Compositional matrix adjust. | **306/464(66%)** | 364/464(78%) | 30/464(6%) |

```
Query  27   TLFKTQHRRYTSFAEDLEYNDFSSGTVGFGQKVSASISRYGDLVSDIMLEVSLPAIEAPL  86
            TLFKT HRR+TSFAEDLE NDFS+GTVGFGQKVSA+ISRYGDLV+D+ +E+ LP IEA
Sbjct  2    TLFKTVHRRFTSFAEDLEENDFSAGTVGFGQKVSATISRYGDLVTDMFMEIHLPPIEA--  59
Query  87   EAAYITXIIPARTGLTATDASGVALAAGVTTGFSAYWVNAIGFALISEVQIEIGGTEIDT  146
                          +  TDASG  +A       +AYWVNAIGFALISEVQIEIGGTE+D
Sbjct  60   -------------AVNVTDASGNTVADADK---AAYWVNAIGFALISEVQIEIGGTEVDI  103
Query  147  LYPEWMYFWEEMTQRPGARLGEQIGKFSYSDTVEADMVEFASQDRTLFVPLQFWFNKYFM  206
            LYPEWM+FWEEMTQRPGARLGEQIGKF+YS  VE DM+EFA Q RTL+VPL FWFNKYFM
Sbjct  104  LYPEWMFFWEEMTQRPGARLGEQIGKFTYSADVEEDMIEFAQQARTLYVPLPFWFNKYFM  163
Query  207  EHGLSIPLIALSYHEIRVRVTFRSIAECTVCTFTDASLNGGGGVNDIEQDIETLVSGLVP  266
            E GLSIPLIAL+YHEI+V+VTFR ++EC    +       G        +   L  G VP
Sbjct  164  EAGLSIPLIALTYHEIKVKVTFRPLSECCCVVYRTEDPTHG--------EYYALAEGKVP  215
Query  267  VNSNTASTLVSADLEARLLVSYVYLDAEERNAFSSVEHEYLITTVQRAQHNITSANATSD  326
            VNS + STLVS+D++A+LLVSYVYLD +ER+AF++ EH YLITT QR  H ITSA ++SD
Sbjct  216  VNSTSGSTLVSSDMDAKLLVSYVYLDKKERDAFATTEHTYLITTTQRQMHAITSAGSSSD  275
Query  327  QVKIFFNHPSNCLTWMIRPSEYLTGAARRRYSVGHKDMFDFSLASGSAAVGLPFGDVTDA  386
            Q+K++FNHPSNCLTW +RP++++T   RRRYSVGH D+FD+SL S S      +GDVTD
Sbjct  276  QIKLYFNHPSNCLTWFVRPTDWMTN--RRRYSVGHMDLFDYSLKSTSDVS--VWGDVTDP  331
Query  387  TKSASLTLNGHSRWPTDLPGLFFRQTVPVMKWTNASDGYMYVFNFSARGGAWQPTSTLNM  446
             K+ASLTLNGHSR+P ++PGLFFRQT P+MKW N S+GYMYVF+FS +GG+WQPTSTLNM
Sbjct  332  VKTASLTLNGHSRFPDNMPGLFFRQTQPIMKWPNCSNGYMYVFSFSLQGGSWQPTSTLNM  391
Query  447  SRIDHVQLELKYGASMPVSDVLIFAESYNLFIVKEGMGGIRYSN  490
            SRIDHVQLELKYG S+P S+V +FAESYNL +VKEGMGG+RYSN
Sbjct  392  SRIDHVQLELKYGTSIPTSNVFVFAESYNLLVVKEGMGGVRYSN  435
```
