## Supplemental File S2-Aev-NL1 sequences for "Discovery of novel RNA viruses in commercially relevant seaweeds *Alaria esculenta* and *Saccharina latissima*"

### Supplemental File S2, *Alaria esculenta* RNA virus-1 (Aev-NL1) RNA and protein sequences

START codon

STOP codon

End sequence similarity

#### RNA1

**Size	2,420 nt**

**Highest match	x**

**Percentage Coverage	x**

**Percentage Identity	x**

**>Alaria_esculenta-virus-NL1-RNA1**

ATTCCATATGACTTCTTATAGTCTTAAACCTGGAGAGAGGGAAAACCTTTCTCTCCCCCAGCCCTCTGGCTTCCCAGACGGGCTGTTGAACACTCTTCGTAGGACTCTACGTTCGAGTAGTGGTTCCTTGCTTGGCGGCAACCACTTTGTAGGAAATCTTCCGCGCAGACCGTGCGAAGGACATGTCGAGGAGCTTCGTAAGCAGCTCCCGGACTTTAGATACAGAAGCCTCAAGAAATTCAAGGCAGTCTGTGAAGAGAGATGTGGCGTTTGTGCCACATCTTTAGTATCATGTGCGGCATACTTAAAGGTGCACGCTCATATACACAAATGTCCAAAGTTTTGCATGCATTTGTTTAAAAACTTCACTCTTTATAAGGAGCAAAAGATATACGATGCTTGCCAATCAATCAAATTCGCGGCATCCAGTTTAGCTCAAGGAGGTTTTAATAACCTCGCCGAGAAGAAATTAAGTGAGATACCTTTCGTATCTCAATTAGAAGATGAGGTAGTGAGCCACAATGCGTGGCCCAACCTTATTGCTTTCCCGCTCTGGAATATCGAGCCAGGGATCGACAAAGAATCTAAGTTTGATCTTAGTTTCTTACCTCCACCGACACAAAAACAGTGTAATGGATTTCGGAAGAAATTCCGAAGATATCTCGATAAGCATATGCCCGAGATAGTCGAAACTCTTTCTTACACAGAGTGTATGAAAGTTGGCCCGAACAAATTTTATGATGACGGCGAGATTAGAAAGGATAGCGAAACCGCTATCAATTCAGATGGACCATTCTTATATCAGTCCTTCATGACAGGGCCACTCTCTGTACGAGAAGTGTGGCTCCCGACGAAAGCCTTTAAGGCTTCGTCAACGTGGTGGCATCGCGTTGCGGAGCAATTGCTCCACAAACGCCCCCACTTAATACTCTCGCAAGACCCTGAGGCAGCTGCGAGAACAGTACGGAAAAGATTCAAACCGTGCAAATCTATTGACTTAAAAGGCTCAGGCCTTCAGTTTCCGATCGAGTACATAATTATAGTCCTCGAACAGTTAGCGGAACTCTTTCCAGAAATGGAAGAGCGCAAGGACATCGCGGTAGATCTTCTACAGCGTATGTCCATCTTTAAGGATAAGAAATTCCATATCCCGACTCGTGGAGTTGGACTCGGATATTTTACAAATATCAAGGTCATGGTCATAGATTGTCTTCTTGAAGACTACTATGTTATTGCATCCTTCGACGACGATATGCTTGTCAAGGATAATCAGTATAACGCTTGTATTAAGCGGCTGAAATCTTACGGCTTTCTTATTAATGAAGAAAAGTCCGGACATCACTGGCCTGTAAACCCGTGGTTCCTTAATGTTGGCATTCTCATGGATGAAGAGACTGTCTTAGGCTACAGTACGTGTAATGCGTACATGGCAGCCGCATTTACTAAACGCTATCATTGGGAGCGTAAAAGTATTCTCCAACAGGTATACCCTGAGGACTCTCATTATATGGCATTCCATTATGAGAAAATATTTGGATATGAATTCTTCAAAGGAGAATCCATTGAACACCCACACAACGGAGGTCTCTGTTGGTGGAGTAAAGAGCTCGGTGGTAAAGACCAGGGAGCTTTCCTTCAGGCACATCTGATGCCTCAGAGGATCTATGACGAATGGAACGGGGAAATACCCTATCCAACGTTCGCAGCCGACACTATATCGAGCGCGGATAGAAAGCAACACCATTTCAAAAGAAAGAGGATGTTCAAAGATAAGACCATGATTTACTCATGGGATTATTATCATCTCCACCCTCGTATGAGGTCCGGAGAGTCCATTAGGTCGCATTCTAGCGATCTTGACGGAATTACGCCTGCTTGGAGAGAAATACTCCTCCTCAAGCATTTCGGGGTTTCACATGGAAGCCTCGAGCAGGATATACCGAAAGCAGTGATACCACTTCTGGCGGATAAGTTCCCTCTCTCGCGAGATCCAATCCAGGCATTTGCCAGGAGCGAGTTTTGCGATTCAGAGAATGAACCACAAGGAATCGTCTGTGATGAAACACGAGAACGATTACTGGCCATTTCCAGGGCTACGAAGGTCGGCCACGAATACTTCTTTCAGAAACGTGAAGAAGTTAATCAGGATTTGGAAATTCCTCCAAAACCTAACCCTACTCCCCAACTCGATAATCTCTTGTTCGGGGATCTCGAATGGTCCGGTCCGGATCTTTCGGGAATTGAAATTCAGGCTCCCTATGAGGAATCGTCTGAATCTGAAGGAGAGGAAATCCTTTCCGACGCTATGTCCGACTGTATGTCGAATATTAGTGAATTCCTCGACGACATCTAATTAGATGTTGTCGATAAGCTATAATCCCAGTTAAGGGTTTTGGCAGATATGTGCTTTACAGTACACA

#### RNA2

**Size	2,040 nt**

**Highest match	x**

**Percentage Coverage	x**

**Percentage Identity	x**

**>Alaria_esculenta-virus-NL1-RNA2**

ATTCCATATGACTTCGTTTAGTCTAAAATTGCCTGGGGGGGAGAAGCTCTCCCAGGCGCTGGCCGAGGTAGCATGCTACCCGGCCAATGTACAAATCTCGCACATAGAGCGCAAGATTACCGAAGATCTGATTCTCTTTTCTTCGGATGAGAACATTGCATCGATTACAGATGCATGTAACTACTGTACCCCTCATATCGAGGGCTCAGTAATCATTGAACCAGCCTTACAAATGAGGCAGTTCGTCGATCGCGTTAGTAAATTTGTCTATCGCGCTCACACTTCGACAACTCTTAAAGAAATAGAAGAGTTGCAATCCAGCATGCAAAATTTAAGTATGTCTGGAAACACTAATGAACAAGAAGATTTCTCGTTCATGATAGTGCCCGGCAAGCGTGGGGCCTACGAGATCCGCAAGTTAATTCTAACTATACCGGATCTTTCAAAATCAGATCAGTCACATAGGACAAATCTGGTTGAACACATCAAGTTCGCTGCGACCTTGTTGAGTATTAATACTGAAGATATCAAAGTTGATAGAGTCAGCTCGTTGGGGACAACATGTCCGTTACCCGCGAAACTAGGAGGTGCAATGACAGAAATGCTGACATTACACAGATCGTCCACTGCTTCTGCAGCAGTAGGACAATTTAAACTGGAGCGCGACCTTAGGGTCACGACCGGAGGTGCTCTCGCCATTTTGGCGCACTTGCACAAAAGATCACCCTTCTACCGTCGAGACGGAAAGGGAAAATACGTCACGTCTGAGTTACTCAAGACGGTCGTGAACAATGCATTTGGGCTCAATGAGTCCAGATGCTCTCAGTTTTCGAAGAGTTTCTTCAAAGCTGTGTTTAGAGCGATCGTGACAAACGACCTCGTTCGCGTACCTTCGAGCTTCAGTAAAAGCGCGAAGGTGACTTTTGATGTCTCCTCACCTGAAGGCATCATGAGGAAGGCGGGTTATACACCGCTCATTCCCGATACTACCAAGATGCTCTTGGTATTAACGACGGTCGACAAACTCGACCATCGGCTCTTCGGGTCTACTGGCGACCACGAAGAAATCGAGGATACTGGGCAAAAGCCTAAGGATCCTCCGGCTACGGAAGTAGCCACATCACCCCCGGCTGATACGCCTAAGGTGACTTTCGCTCCCGGAACATCAGAAGATGATAAAGGGAAGGGCAAGGCGTCCGATTGGACGACTGTCAAGAAAAAGAATCCATCCAAAAAGGATCGGAGGAAAAAGTCTACTGGGGATCTCCCAGTAGGCCCTAAGTCTAATGACAAGCCTCAAGGCTATGTCATGGAGGCGCCGGCAAAACCGGCAGCTAAGAAAAAGGTTCATCTTGAAACTATTTCTAAACTCGACTCTGCGCGTAATACGCACAAAGCATTCGGGGCGGGGGTGAAACTCCTCCTCCCATTCATCGATCCGAGCTCTAAGCTGTCGATGAAAGACCAGCTCAAATACAGTTATAAGAACTGTTCTGAGCAATCTCTTCTCTTTTTTAAGGAGCAGAGAAATTTTGTCGCGAGCGCCGAAAAGACGTACGCGGTTCTTCAGGCTTCTAAGAACCCGAAGAGCAAAGCCACGGCTGAGCATTATGTGCTAGCCAGGAATCGCATGTGCAACTCTTTATTGCCCATGAAGTTTGCGGACCGTACAGGAACCTGTTACAAGTCCTACTCTGAAATCCCTTTGGGGGTTCGGAGATACTTCGAAAAGGCGCTCTCCCGAGAAATTCAGGAGAGACCAAAGAGCGATGATATCATTGAACCATCGCTAAAGAAATCCAGGGTCGACACTGTCGACATGGACGTAACACCGACAACTCCGGATACGGAGATGGAGGTACCAGAACCCCCGTTAGACCTATAAGGTTAAAAGGGGTTGGTCATGAGTTGCTAACTCTGACCATCAGTTTAGTTCCTTTAGGAGCTTGACTATTTATGGAACCTTATGCGTATGACGTATAAGATTCTTGACTGCTCTCGGCATCAGAGAAAAGTGAATATGTACCTTACAGTACACAC

#### Protein ORF1

**Size	781 aa**

**Highest match	WPV08070, RNA-dependent RNA polymerase [Verticillium dahliae ormycovirus 2]**

**Percentage Coverage	81%**

**Percentage Identity	32%**

**>Alaria_esculenta-virus-NL1-ORF1**

MTSYSLKPGERENLSLPQPSGFPDGLLNTLRRTLRSSSGSLLGGNHFVGNLPRRPCEGHVEELRKQLPDFRYRSLKKFKAVCEERCGVCATSLVSCAAYLKVHAHIHKCPKFCMHLFKNFTLYKEQKIYDACQSIKFAASSLAQGGFNNLAEKKLSEIPFVSQLEDEVVSHNAWPNLIAFPLWNIEPGIDKESKFDLSFLPPPTQKQCNGFRKKFRRYLDKHMPEIVETLSYTECMKVGPNKFYDDGEIRKDSETAINSDGPFLYQSFMTGPLSVREVWLPTKAFKASSTWWHRVAEQLLHKRPHLILSQDPEAAARTVRKRFKPCKSIDLKGSGLQFPIEYIIIVLEQLAELFPEMEERKDIAVDLLQRMSIFKDKKFHIPTRGVGLGYFTNIKVMVIDCLLEDYYVIASFDDDMLVKDNQYNACIKRLKSYGFLINEEKSGHHWPVNPWFLNVGILMDEETVLGYSTCNAYMAAAFTKRYHWERKSILQQVYPEDSHYMAFHYEKIFGYEFFKGESIEHPHNGGLCWWSKELGGKDQGAFLQAHLMPQRIYDEWNGEIPYPTFAADTISSADRKQHHFKRKRMFKDKTMIYSWDYYHLHPRMRSGESIRSHSSDLDGITPAWREILLLKHFGVSHGSLEQDIPKAVIPLLADKFPLSRDPIQAFARSEFCDSENEPQGIVCDETRERLLAISRATKVGHEYFFQKREEVNQDLEIPPKPNPTPQLDNLLFGDLEWSGPDLSGIEIQAPYEESSESEGEEILSDAMSDCMSNISEFLDDI

#### Protein ORF2

**Size	625 aa**

**Highest match	USW07205, hypothetical protein [Erysiphe lesion-associated ormycovirus 3]**

**Percentage Coverage	65%**

**Percentage Identity	36%**

**>Alaria_esculenta-virus-NL1-RNA1-ORF2**

MTSFSLKLPGGEKLSQALAEVACYPANVQISHIERKITEDLILFSSDENIASITDACNYCTPHIEGSVIIEPALQMRQFVDRVSKFVYRAHTSTTLKEIEELQSSMQNLSMSGNTNEQEDFSFMIVPGKRGAYEIRKLILTIPDLSKSDQSHRTNLVEHIKFAATLLSINTEDIKVDRVSSLGTTCPLPAKLGGAMTEMLTLHRSSTASAAVGQFKLERDLRVTTGGALAILAHLHKRSPFYRRDGKGKYVTSELLKTVVNNAFGLNESRCSQFSKSFFKAVFRAIVTNDLVRVPSSFSKSAKVTFDVSSPEGIMRKAGYTPLIPDTTKMLLVLTTVDKLDHRLFGSTGDHEEIEDTGQKPKDPPATEVATSPPADTPKVTFAPGTSEDDKGKGKASDWTTVKKKNPSKKDRRKKSTGDLPVGPKSNDKPQGYVMEAPAKPAAKKKVHLETISKLDSARNTHKAFGAGVKLLLPFIDPSSKLSMKDQLKYSYKNCSEQSLLFFKEQRNFVASAEKTYAVLQASKNPKSKATAEHYVLARNRMCNSLLPMKFADRTGTCYKSYSEIPLGVRRYFEKALSREIQERPKSDDIIEPSLKKSRVDTVDMDVTPTTPDTEMEVPEPPLDL

#### **RNA-dependent RNA polymerase [Verticillium dahliae ormycovirus 2]**

**Sequence ID: 	WPV08070.1**

**Size: 	793 aa**

**>WPV08070.1 MAG: RNA-dependent RNA polymerase [Verticillium dahliae ormycovirus 2]**

MASYSLNVSLLTKKSEVGISYRGNLLAIKQYRRTLLKQRHPLIGGVVKTIPIDKNHLGCIKVGQELFTNF

VYRELFKWKKALKDELVCKNCLTHLLITAAYLKCHTADYPHTTPLCIWIMKNSLLPEGQEFYASYNLRRA

KIFDLTQNKMELGDLADDIIYKIEKTCLKQDAKTKMVTRELPSPKAFPSFNLDKESKMWLCNEKPPPEEC

ILSVKEKIRKVILKYGPKEITIPAPEAVKSLGPSLYSDGHVPRCDFEKPEITWKYSWTYQKFKTDAQTER

EIWLPPKSYKMCSSWWHFYVEPICKKIPWLVSNDTMKEVRLNLHKRFKPCKSIDLKGFGLQFPHEYILGC

MDVINEVYPCEEAEEYRSSVANMLRILSVKMDDGTHVKPIRGVGLGYFSNIMSLVVASLLEEFDVVQMFN

DDILIPSDHYEKGIKKLTDHKFIINEKKSGHMWHKVPMFANVSMARNGTLMYYEVQGLKAAVFLKKYHYE

RKSIHIAAPWVYRWKQTFHYERIFGYEVRRLESQDHPSDLGLTPSAPFRTGWVKGGLLRKYKSPKDGLSE

EERRIWSISFPWKTPKDDNFRIAREKAIQKYKDHIWYTEYDEYLNPRIEDKVDGKTMNPDFFLAGYSLPR

WADLQSVLFNHETTGRTTMGTYPKRAAYYMLNYLLSDNPIHSWISGGYDVISPFYRIPGVSNTILMLYDS

LKKSNRLSSKHVLKRCPENEPVYLSAGSGIEFMKNIMNQDYAKLENFDVGEIISYESDEDESQIIDLDDD

LDIDYPDEDGQSSAGDFDAEELW

#### Protein similarity

| **Alignment statistics for match #1** | | | | | |
| --- | --- | --- | --- | --- | --- |
| Score | Expect | Method | Identities | Positives | Gaps |
| 269 bits(688) | 2e-82 | Compositional matrix adjust. | 213/665(32%) | 328/665(49%) | 44/665(6%) |

```
Query  27   LNTLRRTLRSSSGSLLGGNHFVGNLP-RRPCEGHVEELRKQLPDFRYRSLKKFKAVCEER  85
            +   RRTL      L+GG   V  +P  +   G ++  ++   +F YR L K+K   ++
Sbjct  28   IKQYRRTLLKQRHPLIGG--VVKTIPIDKNHLGCIKVGQELFTNFVYRELFKWKKALKDE  85

Query  86   --CGVCATSLVSCAAYLKVHA--HIHKCPKFCMHLFKNFTLYKEQKIYDACQSIKFAASS  141
              C  C T L+  AAYLK H   + H  P  C+ + KN  L + Q+ Y +    +
Sbjct  86   LVCKNCLTHLLITAAYLKCHTADYPHTTP-LCIWIMKNSLLPEGQEFYASYNLRRAKIFD  144

Query  142  LAQGG--FNNLAEKKLSEIPFVSQLED---EVVSHNAWPNLIAFPLWNIEPGIDKESKFD  196
            L Q      +LA+  + +I      +D   ++V+    P+  AFP +N+    DKESK
Sbjct  145  LTQNKMELGDLADDIIYKIEKTCLKQDAKTKMVTREL-PSPKAFPSFNL----DKESKMW  199

Query  197  LSFLPPPTQKQCNGFRKKFRRYLDKHMPEIVETLSYTECMKVGPNKFYDDGEI-RKDSET  255
            L    PP ++     ++K R+ + K+ P+ +   +      +GP+  Y DG + R D E
Sbjct  200  LCNEKPPPEECILSVKEKIRKVILKYGPKEITIPAPEAVKSLGPS-LYSDGHVPRCDFEK  258

Query  256  -AINSDGPFLYQSFMTGPLSVREVWLPTKAFKASSTWWHRVAEQLLHKRPHLILSQDPEA  314
              I     + YQ F T   + RE+WLP K++K  S+WWH   E +  K P L+ +   +
Sbjct  259  PEITWKYSWTYQKFKTDAQTEREIWLPPKSYKMCSSWWHFYVEPICKKIPWLVSNDTMKE  318

Query  315  AARTVRKRFKPCKSIDLKGSGLQFPIEYIIIVLEQLAELFP--EMEERKDIAVDLLQRMS  372
                + KRFKPCKSIDLKG GLQFP EYI+  ++ + E++P  E EE +    ++L+ +S
Sbjct  319  VRLNLHKRFKPCKSIDLKGFGLQFPHEYILGCMDVINEVYPCEEAEEYRSSVANMLRILS  378

Query  373  IFKDKKFHI-PTRGVGLGYFTNIKVMVIDCLLEDYYVIASFDDDMLVKDNQYNACIKRLK  431
            +  D   H+ P RGVGLGYF+NI  +V+  LLE++ V+  F+DD+L+  + Y   IK+L
Sbjct  379  VKMDDGTHVKPIRGVGLGYFSNIMSLVVASLLEEFDVVQMFNDDILIPSDHYEKGIKKLT  438

Query  432  SYGFLINEEKSGHHWPVNPWFLNVGILMDEETVLGYSTCNAYMAAAFTKRYHWERKSILQ  491
             + F+INE+KSGH W   P F NV   M     L Y       AA F K+YH+ERKSI
Sbjct  439  DHKFIINEKKSGHMWHKVPMFANVS--MARNGTLMYYEVQGLKAAVFLKKYHYERKSIHI  496

Query  492  QVYPEDSHYMAFHYEKIFGYEFFKGESIEHPHNGGLCWWSKELGGKDQGAFLQAHLMPQR  551
                       FHYE+IFGYE  + ES +HP + GL   +    G  +G  L+ +  P+
Sbjct  497  AAPWVYRWKQTFHYERIFGYEVRRLESQDHPSDLGLTPSAPFRTGWVKGGLLRKYKSPKD  556

Query  552  IYDE-----WNGEIPYPTFAADTISSADRKQHHFKRKRMFKDKTMIYSWDYYHLHPRMRS  606
               E     W+   P+ T   D    A  K       + +KD      +D Y L+PR+
Sbjct  557  GLSEEERRIWSISFPWKTPKDDNFRIAREKA-----IQKYKDHIWYTEYDEY-LNPRIED  610

Query  607  GESIRSHSSD--LDGIT-PAWREI--LLLKHFGVSHGSLEQDIPKAVIPLLADKFPLSRD  661
                ++ + D  L G + P W ++  +L  H      ++     +A   +L   + LS +
Sbjct  611  KVDGKTMNPDFFLAGYSLPRWADLQSVLFNHETTGRTTMGTYPKRAAYYML--NYLLSDN  668

Query  662  PIQAF  666
            PI ++
Sbjct  669  PIHSW  673
```

#### **RNA-dependent RNA polymerase [Verticillium dahliae ormycovirus 3]**

**Sequence ID: 	USW07205.1**

**Size: 	527 aa**

**>USW07205.1 hypothetical protein [Erysiphe lesion-associated ormycovirus 3]**

MTDEIKNIESDLDMLQFAKSFIKTQLIIKDKDILLDFTKDLATKLSLMKISDGVESHERLVKALDTGPSN

HETVKKPYLVVSGNVLVFRIPSYDTKNTGQEDNLGKIMKWAVTLLNVKNPRVRMVRDKNLGNDIVLPARV

SRMMDMTISALTLPPSEAGERSEFKIGFKANLVELLAAIKLLKKNIGLVQKAPTPKRTKSLTISLDDLKK

SVNGRAGLNEHGMPGYLVGIVKEVFNILTKPNTNILPGNWINSLKQTNGVQSSTAVLYKLGYETIVASPQ

KTLTVVKHRVRDREPKNSSEIKQSVSRTVDGKQQKIFSEVYVTDDKNEPTGISHQEFRLGVCMLLPYIDP

SSSLDMKSQISKDPLSVRNRAVLEFYKRNRRIVDLSNMTYATRSALGKKDSKATVRGYQNARYRTFNECI

KCDFMDAHGNTYARFSDIPKDIRGFLCKLMNRKLNASETEPLDDTEGGEPIPLDEDVEMEPSTAPDEEVE

LQAPTRRTRKSKRPEKHTSQVDDMDVFPKRDKVLKSK

#### Protein similarity

| **Alignment statistics for match #1** | | | | | |
| --- | --- | --- | --- | --- | --- |
| Score | Expect | Method | Identities | Positives | Gaps |
| 55.5 bits(132) | 1e-11 | Compositional matrix adjust. | 67/293(23%) | 122/293(41%) | 18/293(6%) |

```
Query  48   ENIASITDACNYCTPHIEGSVIIEPALQMRQFVDRVSKFVYRAHTSTTLKEIEELQSSMQ  107
            +NI S  D   +    I+  +II+    +  F   ++  +     S  ++  E L   ++
Sbjct  6    KNIESDLDMLQFAKSFIKTQLIIKDKDILLDFTKDLATKLSLMKISDGVESHERL---VK  62

Query  108  NLSMSGNTNEQEDFSFMIVPGKRGAYEIRKLILTIPDLSKSDQSHRTNLVEHIKFAATLL  167
             L    + +E     +++V G         L+  IP     +     NL + +K+A TLL
Sbjct  63   ALDTGPSNHETVKKPYLVVSGN-------VLVFRIPSYDTKNTGQEDNLGKIMKWAVTLL  115

Query  168  SINTEDIKVDRVSSLGTTCPLPAKLGGAM---TEMLTLHRSSTASAAVGQFKLERDLRVT  224
            ++    +++ R  +LG    LPA++   M      LTL  S     +  +   + +L V
Sbjct  116  NVKNPRVRMVRDKNLGNDIVLPARVSRMMDMTISALTLPPSEAGERSEFKIGFKANL-VE  174

Query  225  TGGALAILAH---LHKRSPFYRRDGKGKYVTSELLKTVVNNAFGLNESRCSQFSKSFFKA  281
               A+ +L     L +++P  +R  K   ++ + LK  VN   GLNE     +     K
Sbjct  175  LLAAIKLLKKNIGLVQKAPTPKR-TKSLTISLDDLKKSVNGRAGLNEHGMPGYLVGIVKE  233

Query  282  VFRAIVTNDLVRVPSSFSKSAKVTFDVSSPEGIMRKAGYTPLIPDTTKMLLVL  334
            VF  +   +   +P ++  S K T  V S   ++ K GY  ++    K L V+
Sbjct  234  VFNILTKPNTNILPGNWINSLKQTNGVQSSTAVLYKLGYETIVASPQKTLTVV  286
```

| Score | Expect | Method | Identities | Positives | Gaps |
| --- | --- | --- | --- | --- | --- |
| 79.3 bits(194) | 4e-19 | Compositional matrix adjust. | 44/123(36%) | 64/123(52%) | 1/123(0%) |

```
Query  461  THKAFGAGVKLLLPFIDPSSKLSMKDQLKYSYKNCSEQSLL-FFKEQRNFVASAEKTYAV  519
            +H+ F  GV +LLP+IDPSS L MK Q+     +   +++L F+K  R  V  +  TYA
Sbjct  333  SHQEFRLGVCMLLPYIDPSSSLDMKSQISKDPLSVRNRAVLEFYKRNRRIVDLSNMTYAT  392

Query  520  LQASKNPKSKATAEHYVLARNRMCNSLLPMKFADRTGTCYKSYSEIPLGVRRYFEKALSR  579
              A     SKAT   Y  AR R  N  +   F D  G  Y  +S+IP  +R +  K ++R
Sbjct  393  RSALGKKDSKATVRGYQNARYRTFNECIKCDFMDAHGNTYARFSDIPKDIRGFLCKLMNR  452

Query  580  EIQ  582
            ++
Sbjct  453  KLN  455
```
